# Plk4 triggers autonomous de novo centriole biogenesis and maturation

**DOI:** 10.1101/2020.04.29.068650

**Authors:** Catarina Nabais, Delphine Pessoa, Jorge de-Carvalho, Thomas van Zanten, Paulo Duarte, Satyajit Mayor, Jorge Carneiro, Ivo A. Telley, Mónica Bettencourt-Dias

**Author notes:** equal contribution (Delphine Pessoa, Jorge de-Carvalho). co-lead authors (Ivo A. Telley, Mónica Bettencourt-Dias).

## Abstract

Centrioles form centrosomes and cilia. In most proliferating cells, centrioles assemble through canonical duplication, which is spatially, temporally and numerically regulated by the cell cycle and the presence of mature centrioles. However, in certain cell-types, centrioles assemble de novo, yet by poorly understood mechanisms. Here, we established a controlled system to investigate de novo centriole biogenesis, using *Drosophila melanogaster* egg explants overexpressing Polo-like kinase 4 (Plk4), a trigger for centriole biogenesis. We show that at high Plk4 concentration, centrioles form de novo, mature and duplicate, independently of cell cycle progression and of the presence of other centrioles. Plk4 concentration determines the kinetics of centriole assembly. Moreover, our results suggest Plk4 operates in a switch-like manner to control the onset of de novo centriole formation, and that distinct biochemical kinetics regulate de novo and canonical biogenesis. Finally, we investigated which other factors modulate de novo centriole assembly and reveal that proteins of the pericentriolar matrix (PCM) promote biogenesis, likely by locally concentrating critical components.

## Introduction

> “(…) *the problem which has interested cytologists and embryologists for many years, namely, whether an ordinarily self-duplicating body may, under certain conditions, seem to be created de novo.”* (Dirkson, 1961): On *The presence of centrioles in artificially activated sea urchin eggs*.

It was not long after their discovery in cells in the late 1890’s (by Boveri and van Beneden), that scientists began proposing that centrioles were not always assembled through duplication (Harvey, 1936; Yatsu, 1905). The fascinating discovery that such an elaborate yet fully functional structure can form without a template, raised a variety of questions regarding the regulation of organelle biogenesis, many of which stay pertinent to this date. And while much effort has contributed to our current understanding of the regulation of pro-centriole assembly next to an already mature, mother structure, much less is known regarding the “unguided” de novo centriole formation.

Centrioles are cylindrical microtubule (MT)-based structures that assemble centrosomes and cilia in eukaryotic cells. The animal centrosome is typically composed of two centrioles, surrounded by Pericentriolar Material (PCM), a membrane-less compartment, which contains hundreds of proteins organised within distinct domains, that are responsible for anchoring and nucleating MTs (reviewed in Joukov and Nicolo, 2019). Centriole biogenesis is usually tightly regulated to ensure a correct copy number and to prevent a variety of human diseases (Bettencourt-Dias et al., 2011; Godinho and Pellman, 2014; Godinho et al., 2014; Levine et al., 2018; Marteil et al., 2018; Lopes et al., 2018). In proliferating cells, centriole biogenesis occurs through a canonical pathway synchronous with cell-cycle progression, called centriole duplication. Centrioles begin assembling at G1-S transition, whereby a single procentriole forms at the proximal side of each of the two mother centrioles. During mitosis, centrioles undergo centriole-to-centrosome conversion through the recruitment of Cep135/Bld10, Cep295/Ana1 and Cep152/Asterless (Asl), becoming competent for duplication in the next cell-cycle (Fu et al., 2016; Izquierdo et al., 2014; Wang et al., 2011; Tsuchiya et al., 2016). After mitosis, one centrosome is segregated to each daughter cell. This process entails that the location, timing and number of procentrioles assembled in cycling cells is determined by older/mature centrioles (Banterle and Gönczy, 2017; Breslow and Holland, 2019).

Polo-like kinase 4 (Plk4), also called Sak in fuit flies, is a major player in centriole biogenesis in most animal cells (Bettencourt-Dias et al., 2005; Habedanck et al., 2005; Kleylein-Sohn et al., 2007). Depletion or inhibition of its kinase activity prevents centriole formation, while overexpression leads to the formation of multiple centrioles (Bettencourt-Dias et al., 2005; Habedanck et al., 2005; Wong et al., 2015). Plk4 activity and function is regulated by its concentration, which is known to be very low in human cultured cells (Bauer et al., 2016). Full Plk4 activity is accomplished by trans-autophosphorylation of a conserved T-loop residue within its catalytic domain, which triggers kinase activation through a positive feedback mechanism (Lopes et al., 2015). Other centrosomal proteins also regulate Plk4 activity, such as its substrates STIL/Ana2 and Cep152/Asl (Klebba et al., 2015b; a; Moyer et al., 2015; Zitouni et al., 2016; Mclamarrah et al., 2018; Boese et al., 2018; Aydogan et al., 2019). Moreover, at high concentration, Plk4 self-assembles into nanoscale condensates in *Xenopus* extracts and in human cultured cells, which may also regulate centriole assembly (Montenegro Gouveia et al., 2018; Yamamoto and Kitagawa, 2019; Park et al., 2019).

Centrioles can also form de novo in a variety of cell-types (reviewed in Nabais et al., 2018), but the regulation of this process remains largely unknown. De novo centriole assembly occurs naturally in organisms that lack centrosomes and generate centrioles to nucleate motile cilia, such as land plants that produce ciliated sperm (Renzaglia and Garbary, 2001), several unicellular organisms that alternate between non-flagellated and flagellated life-cycle states and in animal multiciliated cells, where many centrioles are produced at once (reviewed in Nabais et al., 2018). While, in most animals, centrioles are lost during female oogenesis and are provided by the sperm upon fertilisation, as they are needed for embryo development (Rodrigues-martins et al., 2008; Varmark et al., 2007), centrioles form de novo in rodents during early embryogenesis (Gueth-Hallonet et al., 1993; Courtois et al., 2012) and in parthenogenetic insects that can reproduce without fertilisation (Riparbelli et al., 1998; Tram and Sullivan, 2000; Riparbelli and Callaini, 2003; Riparbelli et al., 2005; Ferree et al., 2006). In the latter, multiple centrosomes form spontaneously in the egg at late stages of meiosis, two of which are captured for spindle formation and embryo development, thus replacing the centrioles that are otherwise inherited from the sperm (Tram and Sullivan, 2000).

Centrioles can also form de novo in cells that undergo physical, chemical or genetic perturbations. Proliferating cells are capable of assembling centrioles de novo, but only after their centrosomes have been physically or chemically removed (Khodjakov et al., 2002; La Terra et al., 2005; Uetake et al., 2007; Wong et al., 2015). *Chlamydomonas reinhardtii* carrying a mutated centrin copy has defects in centriole segregation giving rise to progeny without centrioles that, within few generations, reacquire centrioles de novo (Marshall et al., 2001). Although in these cases there is no strict control over the number of centrioles formed, it has been proposed that resident centrioles negatively regulate de novo centriole biogenesis (Marshall et al., 2001), and that such inhibitory effect can be accomplished by having a single centriole in the cell (La Terra et al., 2005; Lambrus et al., 2015).

In *Drosophila* tissue culture cells, evolutionary conserved centriolar components, such as Sas6, Sas4 and Bld10, are critical for both canonical and de novo assembly (Rodrigues-Martins et al., 2007), suggesting that centrioles assembled by both pathways share their core composition but perhaps differ in their triggering. Despite the wide spread circumstances in which centrioles form de novo, the regulation and role of older centrioles on this process have not been addressed. This is in part due to the lack of a controlled model system suitable for high-resolution time-lapse imaging and amenable to experimental perturbations.

In this study, we investigated the spatio-temporal kinetics of de novo centriole assembly, including the effect of pre-assembled centrioles on the biogenesis of new ones, in an egg extract assay to track this process visually, and where PLK4 is upregulated. Plk4 upregulation drives de novo centriole biogenesis in unfertilised *Drosophila melanogaster* eggs (Rodrigues-Martins et al., 2007; Peel et al., 2007). The fly egg is ideal to study centriole assembly since all the proteins necessary for the first centrosome and nuclear cycles are maternally inherited and, in the absence of fertilisation, centrioles are not present. Therefore, centrosomes detected in unfertilised eggs result from de novo assembly and not from duplication from paternally inherited centrioles. Here, we accomplished, for the first time, live-imaging of de novo centriole assembly with high spatial resolution in single-egg cytosolic explants (Telley et al., 2013; de-Carvalho et al., 2018). We show that, at elevated Plk4 concentration, centrioles form de novo and then become competent to duplicate. Both pathways are concurrent as we show that de novo centriole formation occurs independently of pre-existing centrioles. These results demonstrate that in *Drosophila* eggs upon Plk4 overexpression, resident centrioles do not inhibit de novo biogenesis, unlike in human cells and mouse developing embryos. We show that Plk4 modulates the kinetics of centriole assembly in a concentration-dependent manner that is suggestive of a switch-like molecular mechanism occurring in the cytosol. Finally, we find that the PCM, in particular γ-tubulin and Cep152/Asl, strongly regulates de novo biogenesis, suggesting that a local environment of concentrated centriolar and PCM components is required for de novo centriole assembly.

## Results

### An assay to investigate centriole biogenesis live with high spatio-temporal resolution

De novo centriole assembly has been poorly studied in live samples due to the lack of a suitable system where the process can be triggered and documented in a timely manner. Overexpressing Polo-like kinase 4 (Plk4) drives de novo bona fide centriole biogenesis, validated by Electron Microscopy (EM), in unfertilised *Drosophila melanogaster* eggs (Rodrigues-Martins et al., 2007), but the onset of the process and its spatio-temporal kinetics was unknown. Reasons behind this knowledge gap are mostly imaging-related, for the axial depth is optically limited and greatly impaired by the light scattering properties of the egg yolk. Therefore, it is currently impossible to visualise events that take place deep inside the fruit fly egg, which would otherwise be the ideal system to address critical questions concerning centriole biogenesis.

We adapted for this purpose a cell-free assay that overcomes these limitations by generating cell cortex-free micro-scale explants that can be fully imaged, while retaining the native characteristics of the cytoplasm in vivo (Fig. 1A) (Telley et al., 2013; de-Carvalho et al., 2018). Using this assay, we observed de novo centriole biogenesis, at high spatio-temporal resolution (Fig. 1B,C and Suppl. Movie 1). Germline-specific mild overexpression of Plk4 (Suppl. Fig. 1A) triggers the formation of multiple centrioles in cytoplasmic explants, demonstrating that post-meiotic *Drosophila melanogaster* egg extracts are competent for centriole biogenesis (Fig. 1C), and recapitulating observations in eggs which were not extracted, as reported previously (Rodrigues-Martins et al., 2007). Importantly, explants enable to visualize the first steps of de novo biogenesis, which normally occurs deep inside the egg. Moreover, we never observed the elimination of centrosomes during our time-lapse recordings, showing these structures are stable. While it is not possible to accomplish EM validation of centriole structures in cytoplasmic explants since these egg explants are imbedded in halocarbon oil, which is not compatible with sample processing, validation by EM had previously been performed in intact eggs overexpressing Plk4 using the same genetic constructs (Rodrigues-Martins et al., 2007). Given that *Drosophila* egg explants retain and recapitulate fundamental developmental and cytoplasmic properties (Telley et al.,2012,2013), the expectation is that the structures observed here are bona-fide centrioles. Therefore, these extracts offer a powerful assay to investigate the regulation of centriole assembly.

**Figure 1.**
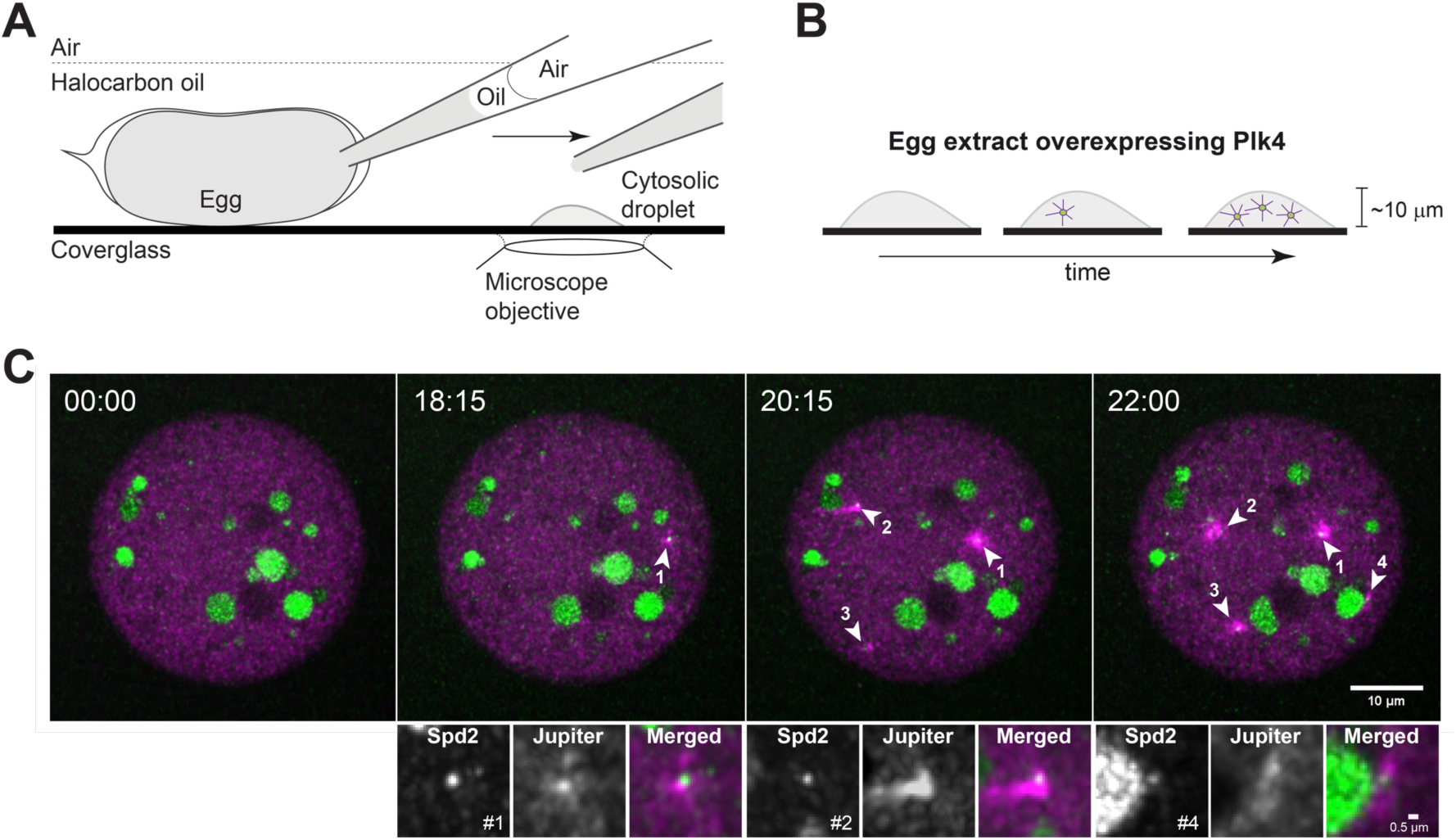
Visualization of centrosome biogenesis in *Drosophila* egg extract. **(A)** *Drosophila* egg extract is prepared by rupturing the membrane and aspirating the cytoplasm with a micropipette. The content is deposited as a droplet on functionalized glass surface. **(B)** Each explant is followed by 3-dimensional time-lapse imaging, documenting centriole formation over time. **(C)** Maximum intensity Z-projections from a time-lapse of a droplet of cytosolic extract isolated from a *Drosophila* egg overexpressing Plk4. Centrioles are absent in the first time point and form de novo throughout the experiment detected as spots (Spd2, in green) associated with a microtubule array (magenta) [arrowheads, numbers indicate the order of birth], reported by the microtubule associated protein Jupiter. The two channels are mostly detected simultaneously, and we have not observed any clear trend of one channel appearing before the other one. The larger green circles are yolk, and the high background is caused by other lipid granules which are highly autofluorescent in the green spectrum, and which cannot be avoided. The insets depict the first centrosomes formed de novo in this time-lapse. The numbers represent their order of appearance. Time is reported as min:sec.

We tested explants of several fluorescent protein fly lines, namely Ana1–dTomato, GFP– Plk4, Asl–mCherry and Spd2–GFP. We chose Spd2–GFP as our routine centrosome reporter because its fluorescence signal was brighter and more photostable across explants than all the others tested, and in our experience this line does not perturb centriole biogenesis (unpublished and de-Carvalho et al., 2020).

### De novo formed centrioles mature and acquire the ability to duplicate in the absence of cell cycle progression

It was previously proposed that in both human cells (La Terra et al., 2005; Lambrus et al., 2015) and *Drosophila* eggs (Rodrigues-Martins et al., 2007), centrioles that form de novo can then duplicate canonically. However, this was never confirmed directly and raises some questions; centriole duplication is thought to depend on centriole maturation, a process called centriole-to-centrosome conversion which requires Ana1 and Asl and is coupled to cell cycle progression (Wang et al., 2011; Izquierdo et al., 2014; Fu et al., 2016; Chang et al., 2016). Centriole duplication is also known to be coupled to cell cycle progression, which does not occur in eggs (Horner et al., 2006; Vardy and Orr-Weaver, 2007; Deneke et al., 2019). Thus, we asked whether centrioles formed de novo mature, recruiting Plk4, Ana1 and Asl, and undergo *bona fide* centriole duplication (Fig. 2 and Suppl. Movies 2A–D). Surprisingly, we observed that recently assembled centrioles can recruit Plk4, the trigger of biogenesis (Fig. 2A,B top panel), Ana1 (Fig. 2C,D top panel) and Asl (Fig. 2E,F top panel) in the absence of cell cycle progression.

**Figure 2.**
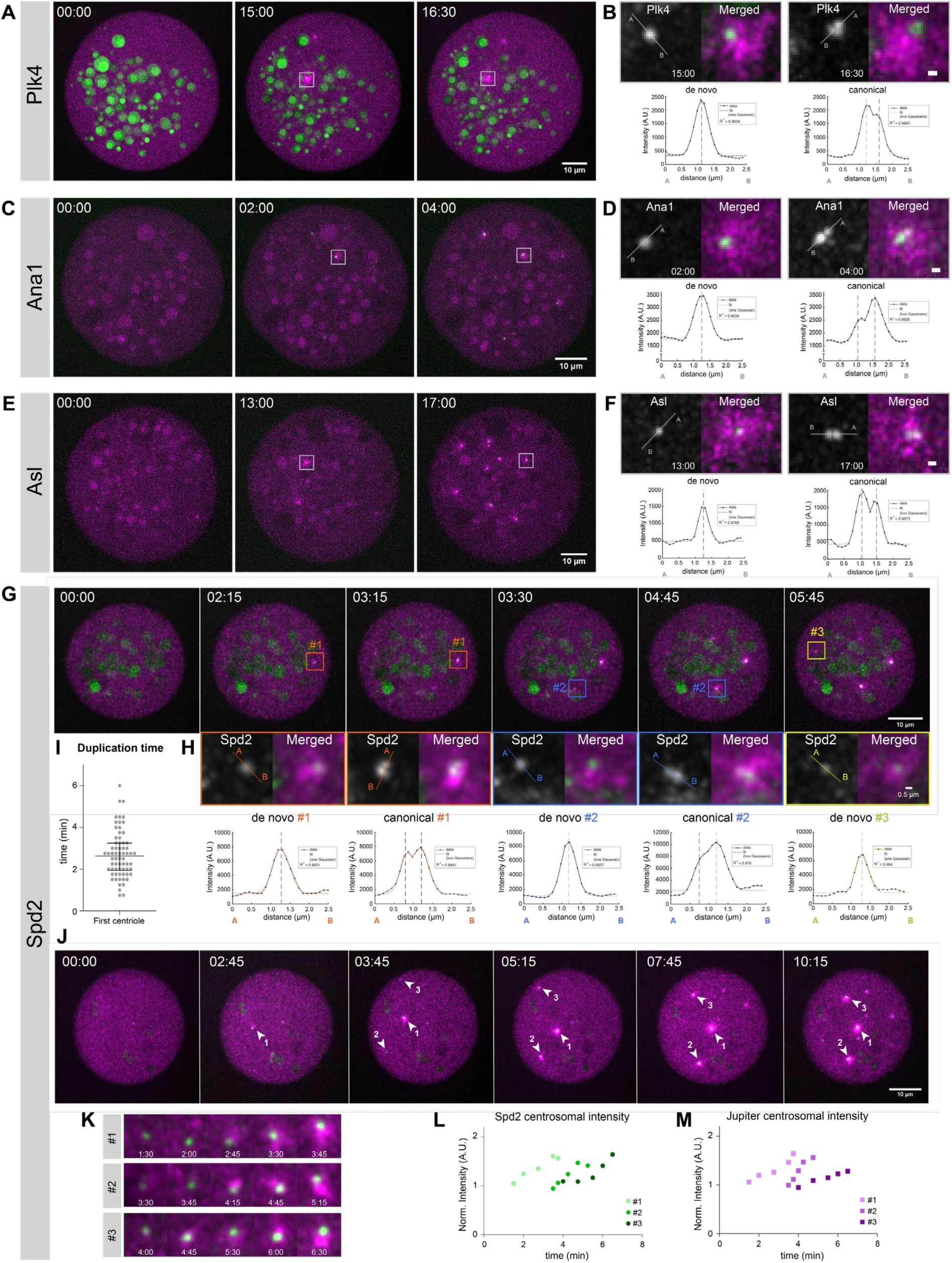
Centrioles assemble de novo, mature and duplicate within the same explants, in the absence of cell-cycle progression. Images show maximum intensity Z-projections from time-lapse movies of cytoplasmic explants extracted from non-cycling unfertilised eggs overexpressing Plk4. Newly assembled centrosomes load Plk4 **(A, B)**, Ana1 **(C, D)**, Asterless (Asl) **(E, F)** and Spd2 **(G, H, J)** shown in green and nucleate microtubules as reported by the microtubule-associated protein Jupiter (magenta). The larger green blobs result from yolk autofluorescence, highly noticeable in the Plk4 and Spd2 panels. **(B, D, F)** Centrioles formed de novo also duplicate, which was inferred from changes in the intensity profile across the centrosomal signal (bottom plots); from a symmetrical Gaussian curve to a Gaussian mixture, suggesting the presence of more than one diffraction-limited structure (centriole). A uni- or bimodal Gaussian distribution was fitted to each “de novo” and “canonical” intensity profiles, respectively (dashed lines represent modes from fit). The coefficient of determination (R^2^) is presented for each fit. Scale bars, 0.5 μm. **(G)** Centrioles form de novo and canonically over time, therefore both biogenesis pathways co-occur. Centriole duplication was inferred from the change in the intensity profile across the Spd2 signal **(H**, bottom plots). Uni- or bimodal Gaussian fitting as in **B–F**. Colors represent one centrosome that first assembled de novo and later duplicated. **(I)** The duplication time depicted in the graph is the time elapsed between the documentation of the first centriole formed de novo (unimodal density) and the detection of a centriole pair (bimodal density). The horizontal line and error bars represent the median and interquartile range (N=66 explants from different eggs). **(K)** Insets of the first three centrosomes formed de novo in time-lapse **(J)** and their corresponding normalised and bleach-corrected intensity of Spd2 **(L)** and Jupiter reporting microtubules **(M)**, plotted over time. Time is reported in min:sec.

Next, we investigated whether centrioles that formed de novo also duplicate, as predicted by their ability to mature and recruit Plk4. In our assay, single centrioles are first detected as radially symmetrical intensity spots with Gaussian intensity profile (Fig. 2B,D,F,G). Over time, a single Plk4, Ana1, Asl and Spd2 Gaussian intensity profile can evolve into a mixture of at least two Gaussian distributions (Fig. 2B,D,F,H), consistent with the presence of more than one centriole and canonical duplication. We next took advantage of 3D-Structured Illumination Microscopy imaging (3D-SIM), which has approximately twice the spatial resolution of confocal microscopy. Spd2–GFP forms a ring at the centre of the microtubule aster, with an inner diameter of about 230–320 nm when viewed in cross section (Suppl. Fig. 1B, insets). Previous studies have demonstrated that Spd2 also forms toroids at the centrosome in *Drosophila* syncytial embryos, whereby Spd2 projections extend from a central hollow structure, which presumably contains a single centriole (Conduit et al., 2015). In our experiments, smaller structures form adjacent to older centrioles which previously formed de novo, further supporting the onset of canonical duplication in this system (Suppl. Fig. 1B, insets). The fact that duplication is a property which has only been observed for centrioles, and not other MTOCS, such as condensates (Montenegro Gouveia et al., 2018), strongly supports our findings that the structures observed here are bona-fide centrioles structurally and functionally. Importantly, in 97% (66/68) of our time-lapse recordings captured by confocal microscopy, we observed the duplication of the first centriole within 2 to 3 minutes after its de novo assembly (Fig. 2I, scatter-plot).

We next asked whether centrioles are fully converting to centrosomes, maturing also in their ability to nucleate MTs. Indeed we observed that as they age, centrioles continue incorporating centrosomal proteins and increase their MTOC capacity, which is reported by the intensity of the microtubule-associated protein Jupiter (Fig. 2J–M).

### Centriole-mediated regulation of centriole biogenesis

Interpretation of earlier experiments led to the model that existing centrioles play a dominant role in centriole assembly and negatively regulate de novo centriole biogenesis (Marshall et al., 2001; La Terra et al., 2005; Uetake et al., 2007; Lambrus et al., 2015). Whether centrioles can release an inhibitory signal is unknown. On the other hand, it has been suggested that centrioles can act as catalysers of centriole biogenesis, by concentrating centriole components and therefore preventing biogenesis elsewhere (Marshall et al., 2001; Lopes et al., 2015). In particular, given that Plk4 activity is regulated by PLK4 trans-autoactivation, it was suggested that its sequestering at the centrosome, would keep PLK4 activity low in the cytoplasm and prevent de novo biogenesis (Lopes et al., 2015).

We asked whether the appearance of the first centriole can prevent further de novo formation. Surprisingly, despite the assembly of centrioles and their duplication, we continue to see de novo formation (timeline in Fig. 2G,H), challenging the view that existing centrioles have a context-independent inhibitory effect in centriole biogenesis. Notably, these subsequent events are too distant from existing centrioles to be mistaken by a centriole that duplicated and moved away within the time of one frame (Suppl. Fig 1C). To further study their occurrence in more detail, we analysed the spatio-temporal kinetics of de novo biogenesis in our assay, by assessing if centrioles impact the place and timing of other de novo events. Once the first centrosome had formed, we assessed if older centrioles affect the biogenesis of others, e.g. by promoting (triggering effect) or repressing (inhibitory effect) the assembly of new ones either globally or in their vicinity (Fig. 3A,E).

**Figure 3.**
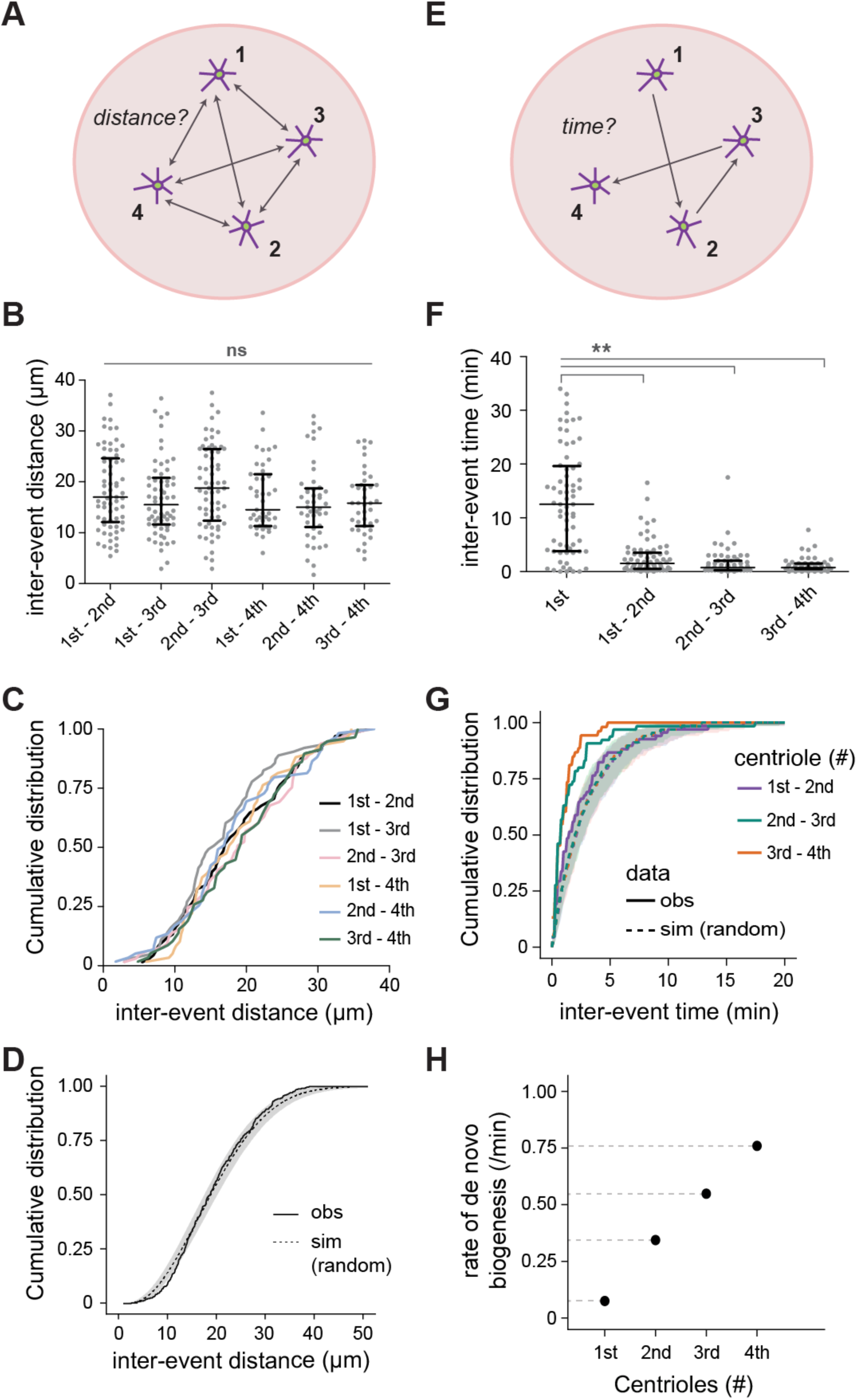
Spatio-temporal kinetics of de novo centriole biogenesis. **(A)** Schematic representation of the experimental data analysis for distances. The first four centrosomes formed de novo in the explants were tracked in 3D using the intensity signal from the Jupiter (MT reporter) channel (first tracking round) and Spd2 (centrosomal reporter) channel (second tracking round) combined. For each of the de novo birth events, an XYZT coordinate matrix was retrieved, from which the inter-event distances were calculated. Experimental N=68 droplets/eggs. **(B)** Scatter plot of observed inter-event distances for all pairwise combinations of the first four de novo biogenesis events. Horizontal lines and error bars represent median and interquartile distance, respectively. **(C)** Cumulative distribution functions of inter-event distance. Distributions were not significantly different (Kruskal-Wallis mean rank test, p-value= 0.467). **(D)** In silico simulations were performed to test if the observed experimental data deviates from a theoretical scenario in which all four birth events occurred at independent and identically distributed random positions with a uniform probability density distribution, within explants with similar geometry as in the experiments. Four random events were obtained in 100 simulations of 68 droplets. The graph depicts the median CDF of all experimentally observed (*obs*, solid line) and all simulated (*sim*, dashed line) inter-events distances, while the grey envelope indicates the 95% Confidence Interval (from quantile 0.025 to 0.975) for the simulated data. The experimental observations do not deviate from random simulations. **(E)** Schematic representation of the experimental data analysis for time. For each of the four de novo birth events, an XYZT coordinate matrix was retrieved, from which the inter-event time were calculated. Experimental N=68 droplets/eggs. **(F)** Scatter plot of observed inter-event time between the first four de novo biogenesis events. Horizontal lines and error bars represent median and interquartile range, respectively. The first event time is significantly different from subsequent inter-event times (Kruskal-Wallis mean rank test, p-value= 0.0047). Note that this 1^st^ event time exhibits high (systematic) variability due to an ill-defined time reference (see Methods) **(G)** Cumulative distribution functions of observed (continuous) and in silico obtained inter-event time (dashed). Simulations were performed to test if the observed experimental data deviates from a theoretical scenario where all four birth events occurred independently at a constant rate within an explant with similar geometry as in the experiments. Four random events were obtained in 100 simulations of 68 droplets. In the simulation, the 1^st^ event rate of birth was approximated to the inter-event time between the first and second events. The graph depicts the median CDF of the experimentally observed (*obs*, continuous line) and simulated (*sim*, dashed line) waiting times between the first and second, second and third and third and fourth events, while the grey envelope indicates the 95% Confidence Interval (from quantile 0.025 to 0.975) for the simulations. The observed and simulated waiting time distributions do not overlap, and differ more as centriole number increases, suggesting that the rate of biogenesis is increasing over time. **(H)** Estimation of the experimental birth rates using Maximum Likelihood (MLE) fitting. An exponential distribution with rate λ>0 was fitted by MLE to the CDF of each observed waiting times. The estimated rate of de novo centriole assembly is represented in the graph as a function of the number of centrioles previously/already present in the volume.

We did not observe a statistical difference in the pairwise inter-event distance between the first four centrioles formed de novo (Kruskal-Wallis mean rank test) (Fig. 3B, Suppl. Fig. 2A, top). However, we noticed that new centrioles form, on average, more than 10 μm away from previous ones, regardless of centriole rank and explant size (Fig. 3B,C), raising the question whether this process is spatially random or if there is any spatial regulation (e.g. an inhibitory effect) imposed by older centrioles on the birth of neighbours. To test these hypotheses, we generated stochastic models with similar geometric constraints as the cytosolic explants, allowing us to compare observed and simulated data. By measuring the simulated inter-event distances between four random events, independent and uniformly distributed within 3-dimensional spaces of similar geometry, we could derive that the experimental observations in explants do not significantly deviate from a random spatial process (Fig. 3D, Suppl. Fig. 2A, bottom).

According to our measurements, older centrioles have only a short-range effect on the biogenesis of new centrioles, in which they promote canonical duplication, which occurs by definition in very close proximity (Fig. 2B,D,F,H). Importantly, older centrioles do not determine the place of de novo assembly elsewhere in the cytosol. Centrioles behave as autonomous entities at the initial stages of de novo assembly on the scale of tens of micrometers. While we cannot exclude that the mild overexpression of Plk4 observed in this extract is sufficient to override inhibitory signals potentially arising from newly born centrioles, our results rather suggest that de novo centriole biogenesis is not affected by the mere presence of other recently assembled centrioles, through either inhibition or activation. Thus, it is possible that biochemical changes at the level of the entire cytosol allow for stochastic de novo centriole formation that is independent of already present centrioles. To obtain further insight we next investigated the temporal kinetics of de novo biogenesis.

### The kinetics of de novo biogenesis

We measured the time for the first four de novo centrioles to appear in the explants (Fig. 3E). We detected, on average, a long lag-phase until the birth of the first event, independently of the centrosome reporter used (data not shown), after which the process seemingly accelerated. The subsequent rate of de novo centriole biogenesis was in the range of one every two minutes (Fig. 3F). We then asked whether these were independent events, such as suggested in our spatial analysis, by running *in silico* experiments.

Assuming independent de novo biogenesis events with a constant rate, computer simulations predict that all inter-event times should follow a similar distribution (Fig. 3G). However, not all of the observed inter-event distributions were within the confidence interval of the simulation. Moreover, the difference was more noticeable at higher number of centrioles (see 2^nd^ to 3^rd^ and 3^rd^ to 4^th^, Fig. 3G). Finally, Maximum Likelihood Estimation of birth rate indicated a linear increase with centriole number (Fig. 3H).

Altogether, our results demonstrate that the rate of de novo centriole formation increases in time and may comprise two distinct phases. In an initial lag phase preceding the formation of the first centriole(s), the probability of centriole assembly is very low. In the subsequent phase events occur in a burst, at a fast pace. The observed kinetics is reminiscent of a bistable process (Tyson and Novak, 2001; Charvin et al., 2009), switching between non-permissive and permissive states of centriole biogenesis. Moreover, given that all centrosomes are retained in the explants once they are assembled, conditions must exist in the cytoplasm to warranty their stability.

Cell-cycle transitions typically show bistability; they rely on accumulation of a signal or activating enzyme, and the moment a critical threshold is crossed the kinetics becomes essentially irreversible and independent of the signal (Tyson and Novak, 2001; Charvin et al., 2009). Not detecting any effect of the first centriole on the spatial distribution of following centrioles (Fig. 3A–D), suggests that concentration and further autoactivation of Plk4 at the first centriole, is unlikely to induce or prevent subsequent de novo events as previously proposed (Lopes et al., 2015). Rather, a cytosolic event as part of Plk4 activation might occur. It is possible that due to its low concentration, Plk4 activation is a rare stochastic event leading to a transition in the biochemical state of the cytosol that promotes the assembly of multiple molecular foci capable of driving centriole biogenesis. Could the observed transition be indeed explained by the accumulation of active Plk4 in the cytosol, or by an unknown event independent of Plk4?

### Plk4 concentration modulates the kinetics of centriole assembly

Full Plk4 activity is accomplished by trans-autophosphorylation of a conserved T-loop residue within its catalytic domain, which triggers kinase activation through a positive feedback mechanism (Fig. 4A, top: from B to A*, and A to A** forms; Lopes et al., 2015). In more detail, as other kinases, newly synthesised Plk4 is autoinhibited, in this case by a cis-interaction between the L1 linker and activation loop (T-Loop). Autoinhibition is relieved upon Plk4 homodimerisation and autophosphorylation of residues within L1 (Klebba et al., 2015a), a process that may be enhanced by Plk4 binding to its substrate Ana2 (Moyer et al., 2015; Zitouni et al., 2016). Moreover, the ability of Plk4 to concentrate in large order oligomers (“condensates”) may further promote its activation and contribute to the onset of centriole biogenesis (Montenegro Gouveia et al., 2018; Leda et al., 2018; Shohei and Kitagawa, 2018; Park et al., 2019). Consequently, the expected kinetics of Plk4 activation may greatly depend on its concentration and thus overcoming a critical concentration threshold (Fig. 4A, bottom). Furthermore, if the observed bistability is a consequence of Plk4 activation in the cytosol, as discussed in the previous section, we expect the rate of biogenesis to be fast once the critical transition has occurred and centrioles start to be formed, provided there is enough activator (Plk4) in the system.

**Figure 4.**
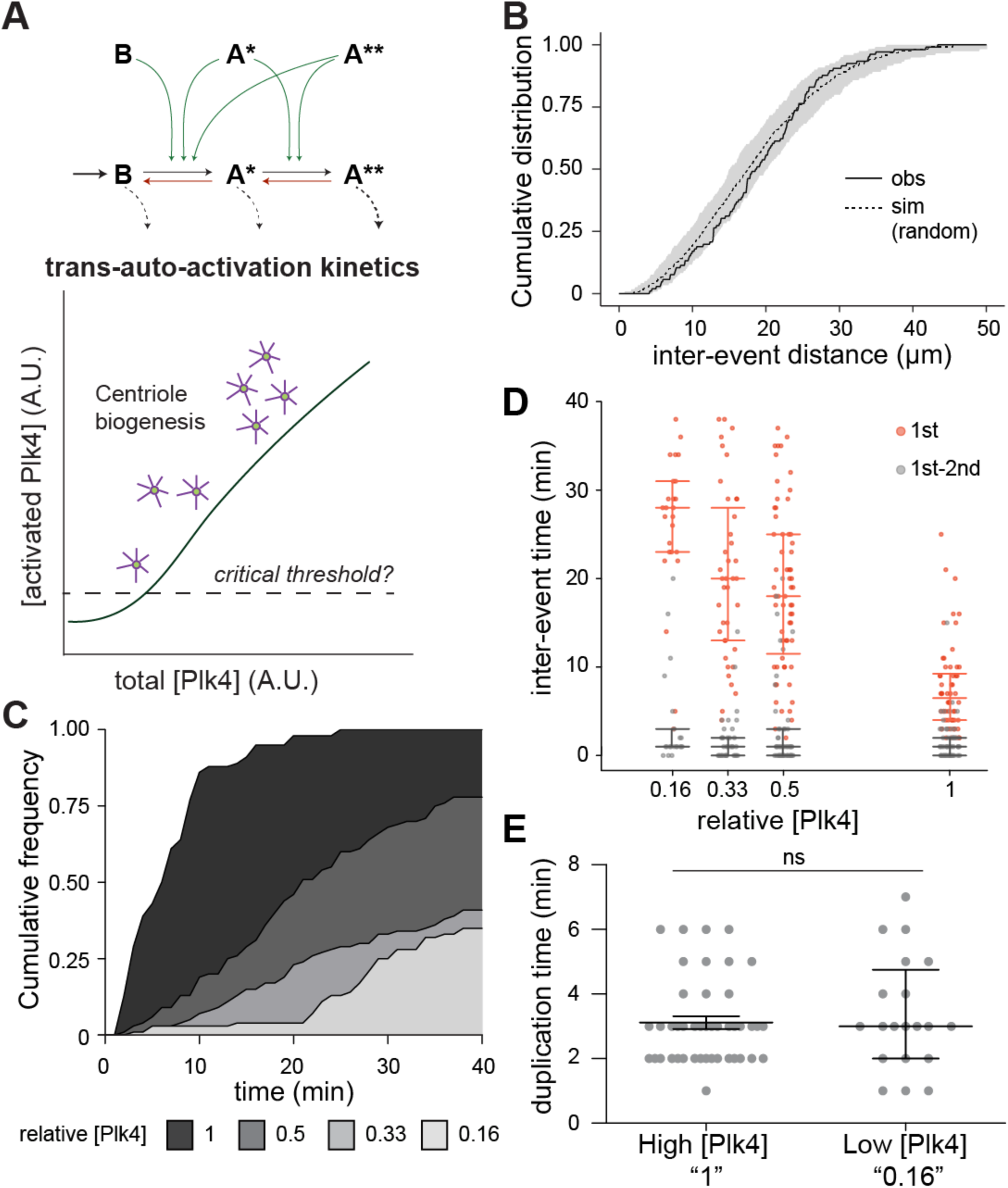
Plk4 concentration modulates the onset of centrosome biogenesis. **(A) Top**. Model of Plk4 autoactivation and dephosphorylation based on data from (Lopes et al. 2015). Plk4 trans-auto-phosphorylates to become fully active, transitioning from an enzyme with basal activity – “B” form – to an activated form phosphorylated on its T-loop residue – “A*” form. Highly phosphorylated Plk4 – “A**” form – is also active but is targeted for degradation (Cunha-Ferreira et al., 2013; Guderian et al., 2010; Holland et al., 2012; Klebba et al., 2013). Dark arrows indicate the forward phosphorylation reaction flux, while red arrows indicate the reverse dephosphorylation flux catalysed by a putative counteracting phosphatase. The leftmost dark arrow marks the synthesised Plk4 that enters the system, while the dashed lines refer to Plk4 degradation. Green arrows depict the Plk4 forms that catalyse the forward flux. **Bottom**. A non-linear balance between phosphorylation and dephosphorylation activities generates a Plk4 critical threshold, as a function of its concentration. Therefore, total concentration (active and inactive) of Plk4 in cells likely affects the timing at which a critical concentration threshold is overcome and triggers centriole assembly (Lopes et al. 2015). **(B)** Inter-event distance at low Plk4 concentration. Cumulative distribution of observed and simulated inter-event distances measured in 3D for the first four centrosomes formed de novo in the explants, at the lowest Plk4 overexpression (“0.16” relative concentration of Plk4). The grey envelope indicates the 95% Confidence Interval (from quantile 0.025 to 0.975) for the simulated data. **(C)** Plk4 titrations were performed by mixing wild-type and Plk4 overexpressing eggs at different ratios. Time of onset of de novo centriole biogenesis is shown as cumulative distribution function for four relative concentrations of Plk4. Lower concentrations delay the initiation of de novo centriole biogenesis, with a large majority of the individual explants not forming centrioles, at lower concentrations, during the observation time. **(D)** Time to the first de novo event, and inter-event time between the first and second de novo events in mixed explants with different concentrations of Plk4. In all dilutions tested, the time for the first event to occur is longer, while the first to second inter-event time is unaffected. Median with interquartile range is presented for N=56, N=62, N=39 and N=25 explants at 1, 0.5, 0.33 and 0.16 relative concentration of Plk4, respectively. **(E)** The duplication time of the first centriole formed de novo is similar at high (“1”) and low Plk4 concentration (“0.16”). Centrioles formed de novo duplicate, on average, 3 min after their biogenesis, at both high (“1”, N = 44 centrioles) and the lowest (“0.16” Plk4 Dilution, N = 20 centrioles) concentrations of Plk4 investigated in Figure 4. The horizontal lines and error bars represent the respective median and interquartile distance. The duplication time is not statistically different between the two conditions (Mann-Whitney test, p-value = 0.59).

To test our hypothesis, we established a titration assay for Plk4 concentration using egg cytoplasm. Wildtype eggs have all the components, except for Plk4, presumably at similar concentrations as Plk4–overexpressing eggs. Thus, mixing egg cytoplasm from these two genetic backgrounds dilutes only Plk4 within a range of full overexpression and endogenous levels. We measured the spatial organization and temporal kinetics of de novo centriole biogenesis for a series of dilutions, in which the larger dilution (0.16), should be close to endogenous levels.

Reassuringly, even at low concentrations of Plk4 close to endogenous levels, de novo biogenesis was indistinguishable from a random spatial process (Fig. 4B, Suppl. Fig. 2B), supporting the idea that de novo centriole biogenesis is not affected by the mere presence of other recently born centrioles, either through inhibition or activation. We then proceeded to investigate the kinetics of de novo biogenesis at different dilutions. We found that all tested dilutions – 0.5, 0.33 and 0.16 relative concentration – delay the onset of de novo centriole assembly (Fig. 4C,D). The delay is dilution dependent; centrosome formation occurs within all explants at the highest Plk4 concentration, saturating within 25 min. Importantly, saturation is not reached within the observation time at lower Plk4 concentrations (Fig. 4C), and the onset of de novo centriole assembly occurs progressively later at larger dilutions i.e. lower concentrations (Fig. 4D), demonstrating that Plk4 concentration is a determinant for the onset of biogenesis.

We next tested whether our observed data is compatible with a model where a biochemical switch occurs in the cytosol, as soon as a threshold of active Plk4 is crossed, promoting centriole biogenesis. To this end, we ran stochastic simulations taking into consideration Plk4 trans-autophosphorylation and dephosphorylation (Fig. 4A). These simulations, largely in agreement with our experimental observations, support the presence of a Plk4 activity threshold (Suppl. Fig. 3A). Strikingly, our results also show that the time from first to second de novo biogenesis event is independent of Plk4 concentration (Fig. 4E, Suppl. Fig. 3B), strongly supporting that Plk4-driven centriole assembly relies on the proposed switch-like molecular process.

In summary, our experiments provide the first evidence *in vivo* that Plk4 triggers de novo centriole biogenesis through a positive feedback loop characterised by a critical threshold of Plk4 concentration at the cytosolic level. Although several lines of evidence indicate that this positive feedback loop could be materialised by Plk4 trans-autophosphorylation, other more indirect molecular mechanisms such as inhibition by Plk4 of a negative regulator of centriole biogenesis cannot be ruled out at this stage. Our results also lead to the hypothesis that in wildtype eggs, where de novo biogenesis does not occur, Plk4 concentration must be very low and undergoes limited oligomerisation in the cytosol, which can prevent auto-activation until the sperm centriole enters the egg and locally concentrates Plk4, leading to its duplication. However, the concentration and the oligomerisation state of Plk4 in the cytoplasm have never been studied in *Drosophila*. Therefore, we decided to investigate these biochemical parameters in the early fly embryo using Fluorescence Correlation Spectroscopy (FCS).

### Plk4 robust regulation under endogenous conditions

FCS is a technique with single molecule sensitivity and, therefore, ideal for quantification of low abundance proteins present at nanomolar to picomolar concentrations inside the cell. FCS measurements of Plk1, also a member of the Polo-like kinase family, revealed distinct diffusion coefficients for Plk1 in the cytoplasm that correlated with its kinase activity during different cell-cycle stages (Mahen et al., 2011). Therefore, we conducted *in vivo* FCS to determine Plk4 concentration, diffusion and oligomerisation in syncytial embryos.

We tagged both Plk4 allelles with a fluorescent reporter by CRISPR to assess its endogenous levels (Suppl. Fig. 4A–C and Suppl. Movie 3). First, we characterized the biophysical properties of the tag mNeonGreen in buffer and injected into control (RFP–β-Tubulin expressing) embryos for reference (Suppl. Fig. 4D–E). Next, we performed FCS in mNeonGreen-Plk4 expressing embryos; we could detect bursts of green fluorescence above background signal (Suppl. Fig. 4F-i). More importantly, the mNeonGreen–Plk4 traces generated clear autocorrelation function (ACF) curves, whereas the background fluorescence measured in RFP–β-Tubulin expressing embryos did not autocorrelate (Suppl. Fig. 4F-ii). For mNeonGreen–Plk4, the normalised ACF were best fitted, with minimal residuals, to a two-component diffusion model, and this fit was corroborated by the distribution obtained from the Maximum Entropy Method (MEM) fit (Fig. 5A, Suppl. Table 4). Two fractions of diffusing mNeonGreen–Plk4 were detected in the cytoplasm: one diffusing at 17.17 µm^2^/s which is similar to the fluorophore mNeonGreen alone (Suppl. Fig. 4E) and another, slower fraction diffusing at 1.49 µm^2^/s (Fig. 5A, Suppl. Table 4). While the first fraction probably refers to Plk4 monomers, the second cannot be explained by homo-oligomerisation alone, suggesting that a fraction of Plk4 may associate with quasi-immobile substrates in the cytosol.

**Figure 5.**
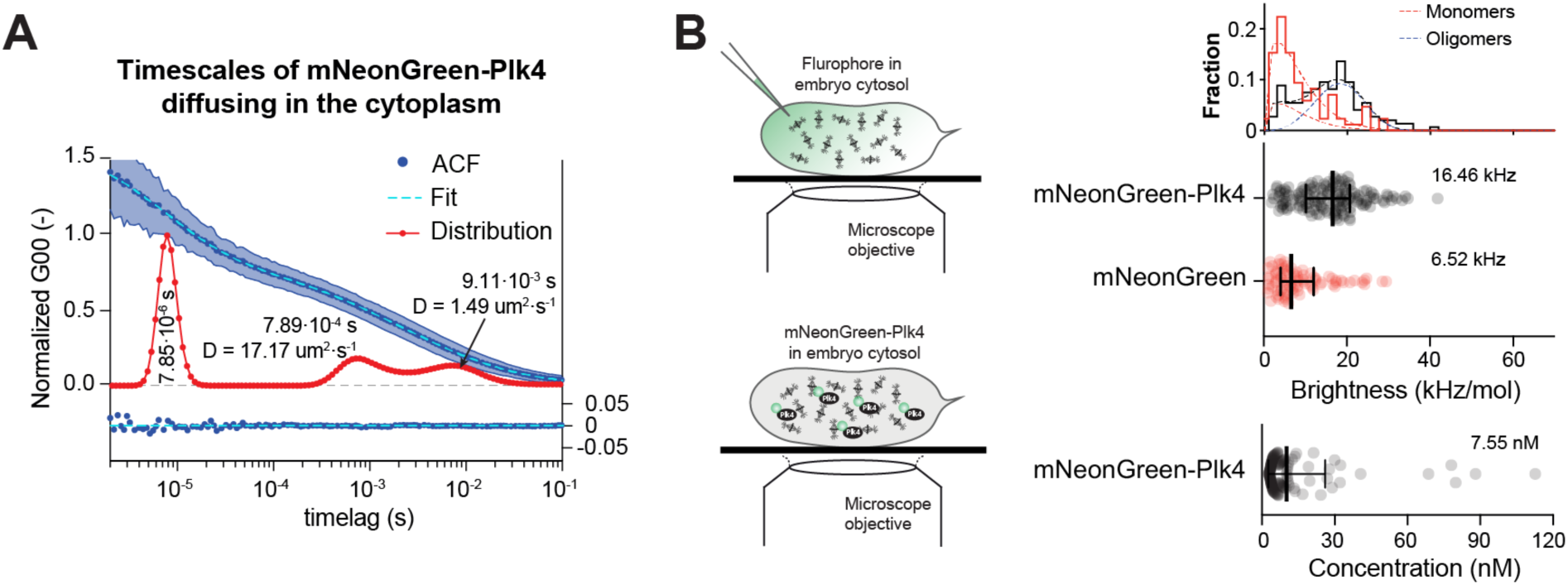
Single-molecule mNeonGreen-Plk4 quantifications in the cytosol of the syncytial fly embryo by Fluorescent Correlation Spectroscopy (FCS). **(A)** Normalised fitted Autocorrelation Function (ACF, “Fit” - light blue dashed line), with standard deviation (shaded area) and Maximum Entropy Method (MEM) distributions (“Distribution” – red line) for mNeonGreen-Plk4 in the cytoplasm. Based on the two fitting methods, three timescales were determined: the fastest timescale peak corresponds to the triplet state of the fluorophore (7.85×10^-6^s); whereas the second and third slower timescales correspond to distinct 3D diffusional mobility of mNeonGreen-Plk4 in the cytoplasm, from which the diffusion coefficients (D) were calculated (fastest fraction: 7.89×10^-4^s, D=17.2 μm^2^/s; slower fraction: 9.11×10^-3^s, D=1.49 μm^2^/s). The residuals from the fitted data (“Fit”) are shown below the graphs. **(B)** Plk4 undergoes limited oligomerisation in the cytosol of the *Drosophila* blastoderm embryo. The mNeonGreen distribution was fitted to a Weibull distribution, which has a peak value of 4100Hz. Next, the mNeonGreen-Plk4 data was fitted with an additional Weibull distribution (one for monomer-like and another for oligomer-like). The second mNeonGreen-Plk4 distribution peaks at 18450 Hz. From this analysis it follows that the overall normalised brightness (intensity per particle, mean ± SD) for mNeonGreen–Plk4 in the cytoplasm is higher than for the single mNeonGreen monomer injected into the cytoplasm at a similar concentration, indicating that Plk4 is present both as a monomer (around 30.1% of its diffusing pool) and as low-order oligomers (69.9% of diffusing mNeonGreen-Plk4 pool).

Next, we calculated the total concentration of mNeonGreen–Plk4 in the cytosol and determined its oligomeric state using the brightness of injected mNeonGreen monomer as calibration (Suppl. Fig. 4E). We confirmed that Plk4 concentration in the cytosol is very low, around 7.55 nM, and an estimate for diffusion in the cytosol suggests coexistence of monomeric and oligomeric form (Fig. 5B). More precisely, 30.1% of diffusing Plk4 is detected as a monomer, while around 69.9% forms low-order oligomers, likely dimers and at most tetramers (Fig. 5B).

Altogether, the FCS results indicate that Plk4 is indeed a very low abundance protein that undergoes limited oligomerisation within the cytosol, in early-developing *Drosophila* embryos. Thus, it is likely that in wildtype eggs, the nanomolar concentration and the low oligomeric presence of Plk4, prevents de novo centriole assembly. Only when centrioles are provided by the sperm, Plk4 can be concentrated, initiating their duplication.

The change in the kinetics of de novo centriole assembly in response to Plk4 concentration allied to the current body of knowledge in the centrosome field, collectively suggest that centriole formation is critically regulated by timely concentration of several centrosomal molecules in one single place (Rale et al., 2018; Takao et al., 2019). But what helps to concentrate these centrosomal molecules? Recent studies suggest that the pericentriole matrix (PCM) may play an important role.

### PCM components promote the early steps of centriole de novo assembly

In *D. melanogaster* cultured cells, co-depletion of the centriolar protein Ana2 and the PCM component D-Pericentrin–like protein (D-Plp) additively impair centriole biogenesis, indicating that two alternative pathways – a centriolar and a PCM-mediated – may be at play (Ito et al., 2019). Moreover, in mouse ependymal cells without centrioles and specialised electron-dense deuterosomes that can feed centriole assembly, a correct number of centrioles can form de novo within Pericentrin rich areas (Mercey et al., 2019b). To test the role of the PCM in de novo centriole assembly, we performed perturbation experiments in *Drosophila* DMEL cultured cells, since it is easier to knock down multiple genes in cultured cells than in the organism. To create an assay for de novo centriole assembly, we depleted centrioles through successive cell divisions in the presence of RNAi against Plk4. As cells proliferate in the absence of centriole duplication, centriole number is progressively reduced. This is followed by a recovery period, without RNAi against Plk4, where Plk4 translation is resumed and centrioles assemble de novo (Rodrigues-Martins et al., 2007).

After RNAi against Plk4, we further depleted PCM components, while allowing Plk4 translation to recover (Fig. 6A), which is sufficient to drive centriole de novo assembly in the mCherry (mCh)-treated control cells (Fig. 6B,C, Suppl. Fig. 5A,B). After 10 days, only 3% of the cells treated with RNAi against Plk4 had centrioles, whereas in the mCherry-treated control about 80% of the cells had at least one centriole, as expected (Fig. 6C) (Rodrigues-Martins et al., 2007). Cells depleted of centrioles were then treated for four days with RNAi against individual – Cnn, Asl, D-Plp or Spd2 – or combinations of PCM molecules necessary for centriole assembly through the canonical duplication pathway – Cnn + Spd2, Cnn + D-Plp, Spd2 + D-Plp or Cnn + Spd2 + D-Plp (Gomez-Ferreria et al., 2007; Conduit et al., 2014; Lerit et al., 2015; Feng et al., 2017; Citron et al., 2018). Additionally, we depleted all four PCM components – Cnn + Asl + D-Plp + Spd2 (referred to as “All PCM”), previously shown to be essential for PCM maintenance (Pimenta-Marques et al., 2016), and the downstream PCM protein, γ-tubulin, which is known to be at the core of MT nucleation across species and contribute for centriole duplication in *C. elegans* embryos and human cells (Dammermann et al., 2004; Kleylein-Sohn et al., 2007).

**Figure 6.**
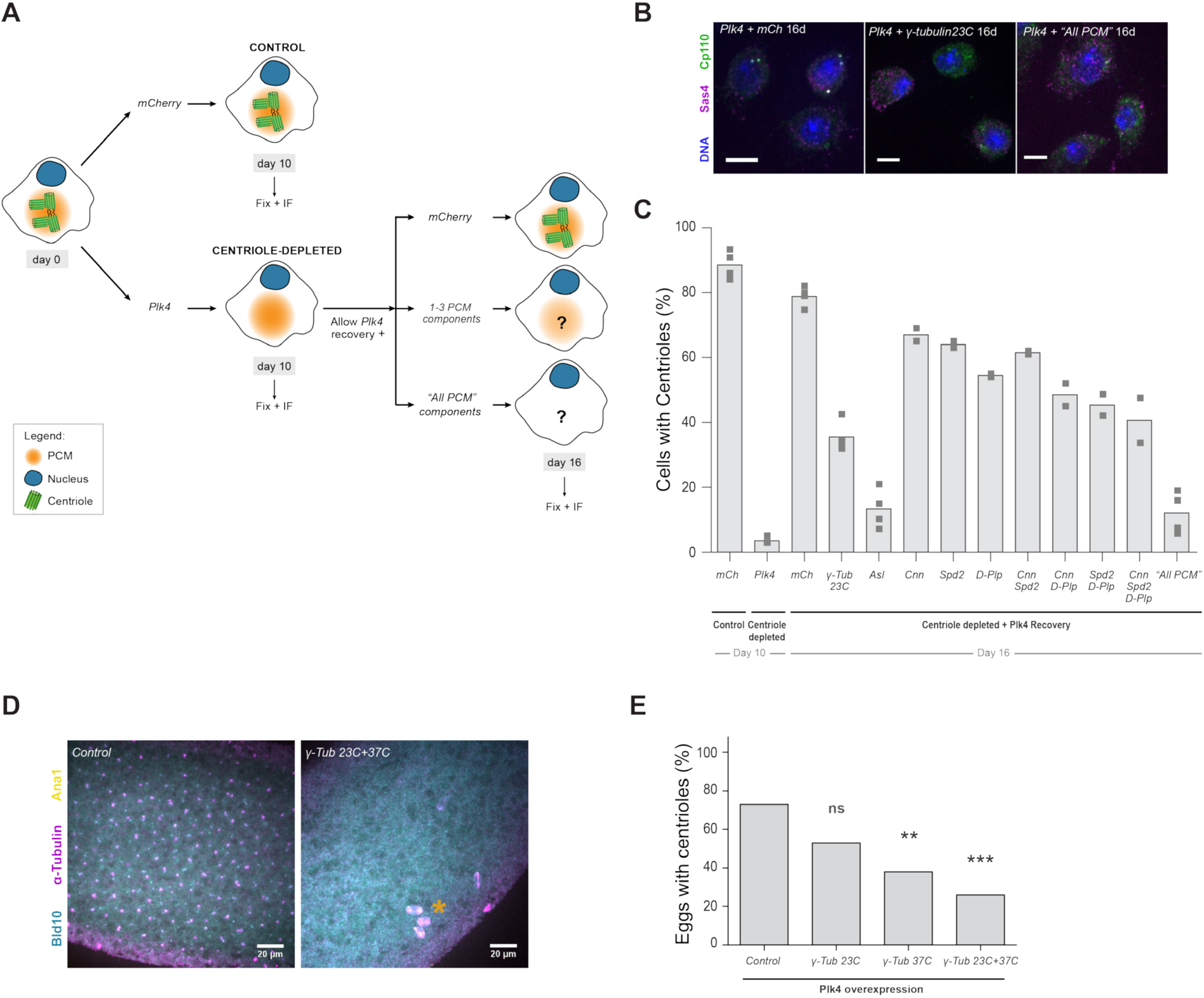
De novo centriole biogenesis is partially impaired in PCM-depleted *Drosophila* cells and eggs. **(A)** DMEL cultured cells were treated with RNAi against Plk4 over the course of 12 days to deplete their centrioles. mCherry (mCh) RNAi was used as negative control. After 10 days, centriole-depleted cells were allowed to recover Plk4 translation while simultaneously treated for four days with RNAi against single PCM components – Cnn, Asl, D-Plp, Spd2 or γ-tubulin 23C – or combinations of these molecules, previously shown to be essential for PCM assembly and/or maintenance. **(B)** Maximum intensity Z-projections of DMEL cells at day 16 treated with RNAi against mCherry (mCh), γ-tubulin 23C or “All PCM”. Cells were stained with antibodies against Sas4 (magenta), Cp110 (green) and DAPI-stained (DNA, blue). Scale-bar = 5 μm. **(C)** Quantification of cells with centrioles after 10 and 16 days of RNAi treatment. Centriole number was scored in > 300 cells per treatment, per experiment. Data is presented as average of two or four independent experiments, depending on the condition. Superscripts ‘*’ denote statistical significance in treatments, where ** and *** indicate p< 0.01 and 0.001 (Pearson’s χ^2^ test and 2-proportions Z-test). **(D)** Maximum intensity Z-projections of unfertilised eggs overexpressing Plk4 alone (Control) or in combination with RNAi against γ-tubulin 23C and 37C together. Eggs were stained with antibodies against Bld10 (cyan), Ana1 (yellow) and tyrosinated α-tubulin (magenta). Yellow asterisk highlights putative meiotic defects, previously described in oocytes from γ-tubulin 37C mutant females (Tavosanis et al., 1997). **(E)** Depletion of γ-tubulin 37C alone or together with γ-tubulin 23C impairs/limits de novo centriole biogenesis in unfertilised eggs overexpressing Plk4. Presence of centrioles was scored in eggs collected from virgin females aged for 4 hours. N=30 eggs (control); N=49 eggs (γ-tubulin 23C); N=47 eggs (γ-tubulin 37C); N=54 eggs (γ-tubulin 23C + 37C). ** and *** indicate p< 0.01 and 0.001, Pearson’s χ^2^ test and 2-proportions Z-test.

While cells treated with control mCherry dsRNA recovered centriole number within 4 days after ceasing Plk4 dsRNA treatment (indicating that centrioles formed de novo), only 10-15% of the cells treated with dsRNA against “All PCM” had centrioles (Fig. 6C), suggesting an important role of the PCM in regulating de novo assembly. From all components of the PCM, Asl, which is known to be both substrate and recruiter of PLk4, generated the strongest phenotype, confirming observations from Dzhindzhev et al., 2010. De novo centriole formation was impaired by γ-tubulin 23C depletion, whereby only about 35% of Plk4 depleted cells recovered a normal centriole number (Fig. 6C, Suppl. Fig. 5B). This result implies that γ-tubulin is important for de novo centriole biogenesis.

We proceeded to validate this observation in vivo and generated fly lines expressing shRNA against γ-tubulin 23C and γ-tubulin 37C (a maternally expressed gene, mostly abundant in early fly development (Tavosanis et al., 1997)), under control of the UASp/Gal4 system. Fertilised eggs laid by females overexpressing the shRNA targeting γ-tubulin 37C do not develop (Suppl. Table 7) and unfertilised eggs display spindle defects similar to those previously shown in oocytes from γ-tubulin 37C mutant females (yellow asterisks in Fig. 6D and in Suppl. Fig. 6A) (Tavosanis et al., 1997), indicating that this RNAi construct is functional. We collected unfertilised eggs expressing RNAi targeting γ-tubulin 23C and/or 37C, while simultaneously overexpressing Plk4, under control of the V32-Gal4 driver. In the control, centrioles form de novo in 73% (22/30) of the eggs overexpressing Plk4 alone (Fig. 6D,E and Suppl. Fig. 6B). On the other hand, in the case of recombinant γ-tubulin 23C + 37C RNAi flies overexpressing Plk4, only 26% (14/54) of their eggs show centrioles, while individual γ-tubulin knock-downs display intermediate phenotypes (Fig. 6E and Suppl. Fig. 6A,B). Therefore, γ - tubulin depletion impairs de novo centriole assembly in vivo too.

## Discussion

De novo centriole assembly is widely documented across the eukaryotic tree of life. Numerous studies reported its incidence and even its relationship with life-history traits in particular groups (Mizukami and Gall, 1966; Aldrich, 1967; Grimes, 1973a; b; Mir et al., 1984; Renzaglia and Garbary, 2001; Idei et al., 2013), but they have not addressed how de novo assembly is regulated in living cells and what the contribution of older centrioles to this process is. Here, we demonstrate that cytosolic explants from post-meiotic *D. melanogaster* eggs overexpressing Plk4 are competent of de novo and canonical centriole biogenesis, offering the opportunity to investigate centriole formation at high spatio-temporal resolution by confocal fluorescence microscopy (Fig. 1). In these explants, Plk4 triggers stochastic formation of multiple centrioles. Our assay allowed us to study several important open questions regarding the regulation of de novo centriole biogenesis.

### Canonical biogenesis is spatially and temporally robust

Our current knowledge supports the need for extrinsic timely cues, provided by the cell cycle, to control biogenesis (Wang et al., 2011; Izquierdo et al., 2014; Fu et al., 2016; Tsuchiya et al., 2016). However, here we observed that centrioles can be formed de novo and undergo time-dependent centriole-to-centrosome conversion, maturation and duplication (Fig. 2), all in the absence of cell cycle transitions (Horner et al., 2006; Vardy and Orr-Weaver, 2007; Deneke et al., 2019). Surprisingly, we also observed that while de novo takes longer to start at low PLK4 overexpressing conditions, duplication time is similar at different mild Plk4 overexpressing concentrations (Fig. 4E). This indicates that, despite the absence of a typical cell-cycle reaction network, canonical biogenesis is both spatially and temporally robust. When it occurs, it occurs always close to an existing centriole and it always takes the same time. Hence, we propose that distinct biochemical reaction networks regulate de novo and canonical biogenesis, with de novo biogenesis being more sensitive to Plk4 concentration.

### A switch-like transition mediated by Plk4 activity in the cytosol promotes de novo biogenesis

In a switch-like process, the system typically undergoes an irreversible transition upon crossing a critical threshold. Evidence supporting a switch-like mechanism operating within the entire cytosol comes from two sources; the measured time of first de novo events being modulated by the concentration of Plk4 while the inter-event time is much less affected (Fig. 4C,D, Suppl. Fig. 3), the spatial distribution falling within random predictions at different Plk4 concentrations (Suppl. Fig. 2). Theoretical modelling and simulations agree with the non-linear kinetics of Plk4 trans-autoactivation in the cytosol shown by (Lopes et al., 2015), suggesting that the burst in biogenesis occurs once a critical activity threshold is overcomed (Fig. 4, Suppl. Fig. 3, also proposed by Lambrus et al. 2015 for the regulation of canonical duplication). Plk4 may also need to oligomerise and form condensates that become stable seeds for centriole assembly (Montenegro Gouveia et al., 2018; Yamamoto and Kitagawa, 2019; Park et al., 2019). Consistent with this we already observe a few oligomeric forms of Plk4 in the cytoplasm at extremely low concentrations of Plk4 (Fig. 5). We suspect that the concentration of active Plk4 increases over time at multiple sites in the cytosol, overcoming the activity of counteracting factors and driving centriole biogenesis almost simultaneously in independent locations in the explants. Once a critical threshold in molecular concentration is locally crossed, Plk4-driven centriole assembly is likely irreversibly catalysed.

### Which factors can help to locally promote centriole formation?

Besides local Plk4 concentration, other factors may play a role in regulating the location of de novo centriole assembly, breaking cytosolic homogeneity. Our experiments support an important role for the PCM in promoting de novo assembly, in particular its component Asl and its downstream effector γ-tubulin. In fact, depletion of γ-tubulin led to a strong reduction in de novo biogenesis, both in vitro and in vivo (Fig. 6, Suppl. Fig. 5,6). We also observed a role for D-PLP, which is enhanced by Cnn and Spd2 perhaps through their function in γ-tubulin recruitment (Fig. 6, Suppl. Fig. 5).

The PCM may generate protein scaffolds in the cytoplasm where centriolar proteins bind with higher affinity, therefore locally concentrating these molecules and forming stable seeds for centriole biogenesis. For example, CEP152/Asl has been shown to recruit and activate PLK4 (Cizmecioglu et al., 2010; Dzhindzhev et al., 2010; Boese et al., 2018), while D-PLP promotes SAS6 recruitment (Ito et al., 2019). Moreover, γ-tubulin promotes MT nucleation, which may attract more components via motor-based transport or through entrapment of proteins with MT-binding capacity, such as Plk4 (Montenegro Gouveia et al., 2018). These manifold properties of the PCM may promote centriole biogenesis within biochemically-confined environments in the cytoplasm. Our results likely provide an explanation why centrioles can form de novo within PCM clouds, both in vertebrate multiciliated cells (Mercey et al., 2019b) and in cycling cells (Khodjakov et al., 2002).

### Do pre-existing centrioles influence the assembly of new ones?

Previous research had suggested that once centrioles form de novo in cells without centrioles, any other events of biogenesis would be “templated”, i.e., follow the canonical pathway (Marshall et al., 2001; La Terra et al., 2005; Uetake et al., 2007; Lambrus et al., 2015). Together, these studies suggest that centrioles negatively regulate the de novo pathway and/or play a dominant role in biogenesis by recruiting the centrosomal components that limit biogenesis. In fly egg explants, we observed that centrioles continue to form de novo long after the first centriole has assembled and duplicated (Fig. 2). Both pathways – de novo formation and canonical duplication – co-occur within the same cytoplasmic compartment, indicating that “older” centrioles and their duplication do not prevent biochemically de novo centriole assembly, even at low Plk4 overexpression (Fig. 4 and Suppl. Fig. 3). Thus, it appears that these pathways are not inherently mutually inhibitory in the fly germline, upon mild Plk4 overexpression. We compared our observations with *in silico* results obtained under similar spatial geometries. These indicate that recently formed centrioles can duplicate but do not influence de novo formation (Fig. 3 and Suppl. Fig. 2). We propose that, given the biochemical bistability, once the cytosol undergoes activation and is permissive for de novo biogenesis, centrioles may form at various locations independently. Our results further suggest that spatio-temporal (local) concentration of Plk4 must be well-regulated in cells to prevent supernumerary centriole formation.

### How do our results fit with naturally occurring biogenesis?

A previous study had estimated 1200–5000 Plk4 molecules per cell in asynchronous human cells, from which around 70 molecules are loaded at the centrosome (Bauer et al., 2016). We generated flies labelled with mNeonGreen at Plk4 genomic loci (Suppl. Fig. 4) and confirmed that the endogenous, diffusing pool of Plk4 is present at very such concentration and undergoes limited self-association in the cytosol in early fly embryos (Fig. 5B). These properties of Plk4 in the cytosol are unfavourable of de novo centriole assembly, ensuring that centrioles form in the right place by canonical biogenesis. Our measurements help building a quantitative framework for the transition of Plk4 molecules from the cytoplasm to the centriolar compartment, which ultimately controls centriole biogenesis.

Finally, we wonder if our findings in *D. melanogaster* relate to the naturally occurring parthenogenetic development in other organisms, including some species of wasps, flies and aphids (Riparbelli et al., 1998; Tram and Sullivan, 2000; Riparbelli and Callaini, 2003; Riparbelli et al., 2005; Ferree et al., 2006). In those cases, multiple functional centrosomes form spontaneously in the egg during meiosis, two of which assemble the first mitotic spindle and trigger normal development. In the case of *D. mercatorum*, the centrosomes that assemble *de novo* can also duplicate and they do so in a cell-cycle dependent manner (Riparbelli and Callaini, 2003). It would be relevant to determine if the burst in centrosome assembly coincides with an increase in global Plk4 concentration or activation in the egg of these species. Just like in our system, a highly variable number of MTOCs are assembled, suggesting the presence of weak control mechanisms against de novo centriole formation in the germline, once the eggs enter meiosis. Further studies aimed at documenting centrosome birth kinetics and their maturation in these natural systems may reveal more about the principles that govern de novo centriole formation and their conservation throughout species evolution.

In oocytes from some parthenogenetic hymenoptera, maternal centrosomes form de novo close to cytoplasmic organelles highly enriched in γ-tubulin, called accessory nuclei (Ferree et al., 2006). Moreover, centrosome ablation in vertebrate CHO cells is followed by accumulation of γ-tubulin and Pericentrin in nuclear-envelope invaginations, hours before bona-fide centrioles are detected (Khodjakov et al., 2002). Interestingly, if treated with nocodazole, acentriolar CHO cells are no longer capable of assembling centrioles de novo (Khodjakov et al., 2002). Therefore, besides substantiating previous studies, our work further suggests that the organisation of PCM-rich foci likely represents the first, essential step for de novo centriole assembly. In the future, it will be important to understand in detail how PCM and MTs contribute to the early onset of centriole formation, and whether their deregulation is associated with supernumerary centrosomes in cancer.

## Materials and Methods

### Fly work and sample preparation

#### *D. melanogaster* stocks and husbandry

All *D. melanogaster* stocks used in this study are listed in Suppl. Table 1. Transgenic mNeonGreen- and mEGFP-Plk4 flies were generated in-house by CRISPR/Cas9-mediated gene editing (Port et al., 2014). Twenty base-pairs guide RNAs (gRNA) targeting the N-terminal region of Plk4, with 5’ BbsI-compatible overhangs, were ordered as single-stranded oligonucleotides (Sigma-Aldrich). The complementary oligonucleotides were annealed, phosphorylated and cloned into BbsI-digested pCFD3-dU6:3gRNA expression plasmid (from Simon Bullock, MRC, Cambridge, UK). Plasmid DNA templates were designed for homologous recombination-mediated integration of the fluorescent proteins mNeonGreen or mEGFP between the 5’UTR and the first coding exon of Plk4. 1-kbp long 5’ and 3’ homology arms were PCR-amplified from genomic DNA isolated from y1,M{nanos-Cas9.P}ZH-2A,w* flies (Suppl. Table 2) (BDSC# 54591). The mNeonGreen and the mEGFP coding sequences were PCR amplified from plasmids (Suppl. Table 2). All fragments were sub-cloned into the pUC19 plasmid (Stratagene) using restriction enzymes: 5’ Homology Arm - NdeI and EcoRI; Fluorescent tag + linker - EcoRI and KpnI; 3’ Homology Arm KpnI and XbaI. Synonymous mutations were performed on the homology arms, removing the protospacer-adjacent motif (PAM) sequence from the donor plasmid to prevent re-targeting. The final donor template for homologous recombination-mediated integration was composed of a fluorescent reporter and a short flexible linker (see sequence in Suppl. Table 2), flanked by 1-kbp homology arms. Two circular plasmids – pCFD3-Plk4_gRNA and mNeonGreen or mEGFP templates – were co-injected into nos-Cas9 embryos (BDSC# 54591 (Port et al., 2014)). Injected flies (F_0_) were crossed to a balancer strain and single-fly crosses were established from their offspring (F_1_). The resulting F_2_ generation was screened for positive integrations by PCR, using primers dmPLK4 5UTR 3 FW and dmPLK4 1exon Rev (Suppl. Table 3). Homozygous mNeonGreen-Plk4 and mEGFP-Plk4 were crossed to pUb–RFP–**β**2-Tubulin flies (gift from Yoshihiro Inoue, (Kitazawa et al., 2014)), establishing stable stocks.

We also generated flies expressing short hairpin RNAs (shRNA) against γ-tubulin 37C and 23C under the UASp promoter and crossed them with the V32-Gal4 (w*; P{maternal-αtubulin4-GAL::VP16}V2H, kindly provided by Daniel St Johnston), at 25°C, to knock-down both genes in the female germline. To generate γ-tubulin 37C and 23C constructs, sense and antisense oligos for each target gene were annealed and cloned into pWALIUM22, using NheI and EcoRI restriction enzyme sites (Suppl. Table 6). Each construct was inserted into different landing sites on the third chromosome by PhiC31 integrase-mediated recombination (Suppl. Table 6). Germline-specific Plk4 overexpression was accomplished by crossing flies carrying the pUASp–Plk4 construct (Rodrigues-Martins 2007) and the V32-Gal4, at 25°C.

Centrosomes were visualised using the following centrosomal reporters: i) pUb-Spd2–GFP (homemade construct, injected at BestGene Inc.); ii) Ana1–tdTomato (gift from Tomer Avidor-Reiss, (Blachon et al., 2008); iii) pUASp–GFP–Plk4 (homemade construct, injected at BestGene Inc.); iv) Asl-mCherry (gift from Jordan Raff, (Conduit et al., 2015)), in combination with either endogenous Jupiter–GFP (BDSC# 6836) or endogenous Jupiter–mCherry (gift from Daniel St Johnston, (Lowe et al., 2014)), as reporters for centrosomal microtubule nucleation.

Flies were maintained at 25°C in vials supplemented with 20 mL of culture medium (8% molasses, 2.2% beet syrup, 8% cornmeal, 1.8% yeast, 1% soy flour, 0.8% agar, 0.8% propionic acid, and 0.08% nipagin).

#### Testing UASp-RNAi lines for developmental lethality

To test for lethality effects of γ-tubulin 37C and γ-tubulin 23C shRNAs alone and recombined, each line was crossed to V32-Gal4 flies. Female progeny carrying the Gal4 and shRNA was crossed to w^1118^ males (10 females x 5 males per vial, 4 independent crosses) and the number of pupae in each vial was counted 9-10 days after each transfer (3 technical repeats were performed). See results in Suppl. Table 7.

#### Embryo/Egg collections

For embryo collections, 3–4 days old female and male flies were transferred to a cage coupled to a small apple juice agar plate (25% apple juice, 2% sucrose, 1.95% agar and 0.1% nipagin), supplemented with fresh yeast paste. Embryos were collected for 1h and aged for half-an-hour. For unfertilised egg collections, around a hundred 5-7 days old virgin females were placed in the cage and 20 minutes collections were performed. At this age, a large majority of eggs did not contain any centrosomes nor did the explants from those eggs. All cages were maintained at 25°C, under 50–60% humidity. The embryos or eggs were dechorionated in 7% Sodium Hypochlorite solution (VWR), washed thoroughly in milliQ water, aligned and immobilised on clean, PLL-functionalised coverslips, using a thin layer of heptane glue. Samples were covered with Voltalef grade H10S oil (Arkema).

#### Preparation of micropipettes and functionalised coverslips

High Precision 22×22 glass coverslips No 1.5 (Marienfeld) were cleaned for 10 min in 3M Sodium Hydroxide, followed by 4 dip-and-drain washes in milliQ water. Next, they were sonicated for 15 min in “Piranha” solution (H_2_SO_4_ and H_2_O_2_ (30% concentrated) mixed at 3:2 ratio), followed by two washes in MilliQ water, once in 96% ethanol and twice again in milliQ water for 5 min each. Coverslips were spin-dried and subsequently treated for 20 minutes with Poly-L-Lysine (PLL) solution 0.01 % (Sigma-Aldrich), followed by multiple dip-drain-washes in MilliQ water. The coverslips were spin-dried and stored in a clean and dry rack.

Glass capillaries (0.75mm inner diameter, 1 mm outer diameter; Sutter Instrument) were forged into glass needles by pulling them on a vertical pipette puller (Narishige PC-10), using a one-step pulling protocol, at about 55% heating power. Using a sharp scalpel, the tip of the capillary was cut, generating micropipettes with 30-35 µm diameter pointed aperture (Telley et al., 2013).

#### Single egg extract preparation

Cytoplasmic extraction from individual unfertilised eggs and explant deposition onto the surface of PLL-coated coverslips was performed on a custom-made micromanipulation setup coupled to an inverted confocal microscope, as previously described in (Telley et al., 2013) and (de-Carvalho et al., 2018). The size of the explants was manually controlled in order to produce explants measuring between 40 - 80 μm in diameter and approximately 10 μm in height, allowing time-lapse imaging of the entire explant volume.

#### Egg immunostaining and imaging

Unfertilised eggs overexpressing Plk4 and knocked down for γ-tubulin were collected from 5–7 days old virgin females for 2h at 25°C, and aged at 25°C for 4 hours. Protocol was conducted according to (Riparbelli and Callaini, 2005). Briefly, aged eggs were rinsed in MilliQ water + 0.1% Tween, dechorionated in 7% Sodium Hypochlorite solution (VWR) and washed extensively with MilliQ water. Using a metal grid, dechorionated eggs were transferred into a scintillation flask containing 50% ice-cold Methanol + 50% Heptane. The vitelline membrane was removed by vigorously shaking the eggs for 3 min. Devitellinised eggs sunk to the bottom of the lower Methanol phase and were then collected into a 1.5 ml eppendorf and fixed for 10 minutes in Methanol at −20°C. Following fixation, the eggs were rehydrated in Methanol:PBS series (70:30%, 50:50% and 30:70%) for 5 min each, washed twice in PBS for 10 min and incubated for 1 hour in D-PBSTB (1x Dulbecco’s PBS, with 0.1% Triton X-100 and 1% BSA), at RT. Primary antibody incubations were performed overnight at 4°C, with the following antibodies: rabbit anti-Bld10 (dilution 1:500; gift from Tim Megraw, The Florida State University, USA); rat anti-tubulin YL1/2 (dilution 1:50; Biorad) and guinea-pig anti-Ana1 (dilution 1:500; kindly provided by Jordan Raff), diluted in D-PBSTB. Eggs were washed extensively in D-PBSTB and incubated with secondary antibodies for 2h at RT - donkey anti-rabbit Alexa 555 (dilution 1:1000; Molecular Probes), goat anti-rat Alexa 488 (dilution 1:1000; Jackson Immunoresearch Laboratories) and donkey anti-guinea pig Alexa 647 (dilution 1:1000; Jackson Immunoresearch Laboratories) in D-PBSTB. Eggs were washed twice in PSB with 0.1% Triton X-100, twice in PBS and mounted onto coverslips in Vectashield mounting media (Vector Laboratories).

Imaging was conducted on a Nikon Eclipse Ti-E microscope equipped with a Yokogawa CSU-X1 Spinning Disk confocal scanner and a piezoelectric stage (Physik Instrumente) with 220 µm travel range. 0.3 µm optical sections were recorded with a EMCCD Photometrics 512 camera using a Plan Fluor 40x 1.30 NA oil immersion objective, controlled with Metamorph 7.5 software. 491 nm, 561 nm and 640 nm laser lines were used to excite the secondary antibodies. Egg counts were tested with a Chi-square test against the null-hypothesis that the outcome is random. Then, each test condition was compared to the control condition with a 2-proportions Z-test under H0 that the proportions of eggs with centrioles are equal versus HA that the proportion in the test is smaller. The significance level for multiple testing was Bonferroni corrected. Significance level was p=0.01.

### Image acquisition, processing and analysis

#### Time-lapse explant imaging on the spinning disk confocal microscope

Centriole formation was followed by time-lapse imaging in explants initially devoid of centrosomes. Explants were imaged at room temperature using a Plan Apo VC 60x 1.2 NA water objective. We have acquired optical sections of 0.45 μm, carefully selecting the total number of stacks (30 to 35 planes) in order to cover the entire volume of each individual explant. The images were recorded with an Andor iXon3 888 EMCCD camera using a Yokogawa CSU-W1 Spinning Disk confocal scanner equipped with a piezoelectric stage (737.2SL, Physik Instrumente), installed on a Nikon Eclipse Ti-E microscope. Dual-colour (488 nm and 561 nm excitation laser lines, at 100ms exposure time), 15 seconds time-lapses were collected with Andor IQ3 software.

#### Image processing

Multi-stack, time-lapse calibrated images were deconvolved with Huygens (Scientific Volume Imaging, The Netherlands) using a Point Spread Function (PSF) automatically calculated from the data set and run in batch mode, for each channel separately. 32-bit deconvolved images were converted to 16-bit and processed using Fiji (NIH (Schindelin et al., 2012)). Maximum intensity Z-projections of both fluorescence emission channels were produced from the time-lapse acquisitions in FIJI, and selected time-frames and insets were further processed with Photoshop CS6 (Adobe). Graphic representations were performed using using GraphPad Prism software (Version 5.0) and the final figures were assembled in Illustrator CS6 (Adobe).

#### Centrosome tracking

Centrosomes were tracked using the Fiji Plug-in TrackMate v3.5.1 (Jaqaman et al., 2008). Centrosomes were identified by the Spd2–GFP localisation at the centre of mass of the microtubule aster. Relying on this criteria, we performed the TrackMate analysis sequentially, starting with the Jupiter-mCherry channel. First, we applied a *3D Gaussian Blur* filter to the images (sigma = 0.7 pixels), facilitating the particle detection on TrackMate using the Laplacian of Gaussian algorithm. The microtubule asters were automatically detected inside spheres of approximately 0.7 µm in radius, adjusting the threshold value for each time-lapse video independently. Next, the first four de novo formed asters were manually tracked from the list of detected particles. A corrected XYZT coordinate matrix of the first de novo events was saved for each video and imported to MatLab R2016b (The MathWorks, Inc.). MatLab was used to build a 3D binary mask with spheres of radius *r* (where r ≥ microtubule aster size), centred at the detected coordinate points. This allowed bypassing incorrect particle detection caused by the large number of green auto-fluorescent yolk particles of intermediate signal intensity, therefore excluding them from the analysis early on. The resulting 3D masks were concatenated into 4D hyperstacks, using the *Bio-Formats importer* plugin in FIJI. The Spd2– GFP images were multiplied by the corresponding 4D binary masks, resulting in a 4D image retaining the pixel intensity values solely within the Jupiter-mCherry ROIs. Next, we used *TrackMate* to detect centrioles within spheres of 0.3 µm radius, combining sub-pixel localisation and a *Median* filter. After detection, the particles were manually tracked. The final centrosome tracks were exported as an Excel MS spreadsheet.

#### Statistics and mathematical modelling

Centrosome tracking data was imported in R version 3.4.1 for further analysis and modelling. The data was analysed in two ways: one aiming at identifying possible spatial constraints in the positioning of the centrioles relative to each other within the explant at the time a centrosome is formed (neglecting time), while the other aimed at understanding temporal constraints (neglecting space). The data was analysed statistically, and simulations were performed in an effort to understand the underlying principles. The details regarding sample size, statistical tests and descriptive statistics are indicated in the respective figure legends and in the main text.

The experimental data was compared to simulated data by calculating the empirical cumulative distributions of each dataset (one experimental and 100 simulated – each consisting of 68 explants) using the function *ecdf* from the *stats* package; and overlapping the median and 95% confidence interval (from the quantiles 0.025 to 0.975) of the simulated datasets’ cumulative distributions with the corresponding empirical distribution from the experimental dataset. Random numbers were generated using the function *runif* from the *stats* library.

For the spatial analysis, each time a new centriole appeared, the 3D pairwise distances between centrioles was calculated and labelled according to appearance relative to prior centrosomes in the explant. This allowed keeping track of event order and, if any spatial effect of existing centrosomes on the appearance of a new centrosome was present, we would be able to detect a difference in their pairwise distances. To test this, the function *kruskal.test* of the *stats* library was used to perform the Kruskal-Wallis rank sum test on the pair-wise distances and labels. To complement this analysis, we decided to compare the distributions of pairwise distances with those expected by a spatially null model whereby centrosomes appear randomly across the available space in the explant. To simulate this null model, sets of random points were simulated in sections of semi-spheres of similar geometry as each of the experimental explants, characterised by height *h* and diameter *d*. To this effect, a height *z* was generated which satisfied 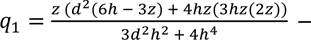 where *q*_1_ was a random number between 0 and 1 – by applying the *optim* function from the *stats* library with the “Brent” method, starting with *z* = 0. This ensured that the *z* coordinate was selected proportionally to the area of the circle it specifies. The two extremes, *z* = 0 and *z* = 1, correspond to the lowest and highest point of the explant, respectively. Subsequently, the coordinates *x* and *y* were generated, within the respective circle at height *z*, by generating a random angle *θ* between 0 and 2*π*, and a random number *q*_2_ between 0 and 1, resulting in *x* = *r* cos(*θ*) and *y* = *r* sin(*θ*), where 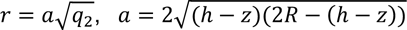 and 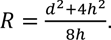. The pairwise distances between simulated points were calculated in the same way as for the experimental data, and the respective empirical cumulative distributions were computed and compared to the experimental empirical distribution, as described above.

For the temporal analysis, the waiting times between centrosome births were calculated from the data and labelled according to which centrosome had just formed. Accounting for a possible change of centrosome birth rate as a function of the number of existing centrosomes, centrosome birth rates were estimated from each of the observed distributions of waiting times by Maximum Likelihood using the *fitdistr* function from the *MASS* library. The experimental data was then compared with a temporal null model whereby centrosomes form at a constant rate in time, irrespective of the existence of other centrosomes and of the volume of the explant. To this effect, random samples of Poisson distributed waiting times were generated using the *rexp* function of the *stats* library, using the rate estimated from the waiting times between the appearance of the first and second centrosomes. The empirical cumulative distributions of these waiting times were compared to those from experimental data, as described above.

The trans-autophosphorylation of Plk4 was modelled following Lopes et al., 2015. Briefly, it is assumed that Plk4 protein is produced with constant source rate s in basal activity form B. The phosphorylation of this B form in the T-loop results in a form A_1_ with higher catalytic activity. The phosphorylation of the A_1_ form the degron converts it to a A_2_ that is targeted for proteasome increasing its degradation rate but that keeps the same catalytic activity. The phosphorylation at the T-loop is catalysed by either low activity B form or the high activities A_1_ and A_2_ forms, while only the later are assumed to phosphorylate the degron of other Pkl4 forms. Both phosphorylation reactions can be reversed by the constant activity of a phosphatase. To keep the stochastic model as simple as possible, we neglected the small first order phosphorylation term in Lopes et al., (2015), corresponding to a putative phosphorylation of Plk4 by other (yet unidentified) kinases. The dynamics of the three Plk4 forms is described by the following set of differential equations:

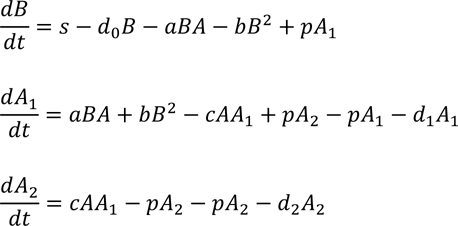

with *A* = *A*_1_ + *A*_2_.

The rate of *de novo* centriole formation in the explant is assumed to be proportional Plk4 activity (*aA*+*bB*) and therefore the probability that an explant has no centrioles F decreases in time according to:

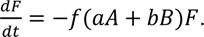

The system of four differential equations was solved numerically using the function *ode* of the package *deSolve* in the software R.

The stochastic solutions for the same set of reactions were obtained by the Gillespie algorithm as implemented in the function *ssa* of the package *GillespieSSA* in R. Each simulation corresponded to an explant where the Plk4 trans-autophosphorylation was simulated independently. The biosynthesis of the first centriole was simulated as a single reaction event that removes a single “precursor” F with a propensity *f*(*aA* + *bB*)*F*. The simulated explant is assumed to form one centriole upon this event.

The model in differential equation and stochastic versions was used to reproduce the temporal evolution of the number of explants containing at least under different concentrations of Plk4. Experimentally four activity levels of Plk4 were obtained by mixing the cytoplasm of eggs overexpressing Plk4 and wildtype, in different proportions with expected activities relative to the overexpressing egg of 1.0, 0.5, 0.33, and 0.12 (Fig. 5B and Suppl. figure 6). The corresponding levels of Plk4 activity were defined in the model through the source parameter *s* = *K*, *K*/2, *K*/3, *K*/6. The value of *K* and the remaining parameters were adjusted by solving the ordinary differential equations for variable *F* and visually comparing *(1-F)* with the experimental time course of the frequencies of explants with at least one centriole (Suppl. figure 6). The adjusted parameters were then used to simulate the stochastic kinetics. The parameter values of the solutions illustrated in Supplemental figure 6 were: *K* = 0.01*Nmin*^−1^*, a* = 1.0/*Nmin*^−1^*, b* = 0.01/*Nmin*^−1^, *c* = 1.0/*Nmin*^−1^*, p* = 0.45*min*^−1^*, d*_0_ = *d*_1_ = 0.01*min*^−1^*, d*_2_ = 0.38*min*^−1^*, f* = 0.34. The value of *N* was set to 2000 molecules for the Gillespie simulations and to the unit in the ordinary differential equations.

#### 3D-Structured Illumination Microscopy

Cytoplasmic explants were imaged with a Plan Apo 60x NA 1.42 oil objective on a GE HealthCare Deltavision OMX system, equipped with two PCO Edge 5.5 sCMOS cameras and 488 nm and 568 nm laserlines. Spherical aberrations were minimised by matching the refractive index of the immersion oil (1.516, in this case) to that of the cytosol, providing the most symmetrical point spread function. 15 seconds, multi-stack time-lapses were acquired, with 0.125 μm Z-steps and 15 frames (three angles and five phases per angle) per Z-section, using AcquireSR (GE Healthcare). Images were reconstructed in Applied Precision’s softWorx software (GE Healthcare) with channel-specific Wiener filters settings (0.003-0.005 for the 488 excitation and 0.005-0.008 for the 568 excitation) and channel-specific optical transfer functions (OTFs). Finally, the reconstructed images were aligned on softWorx and processed using Fiji (NIH, (Schindelin et al., 2012)). Maximum intensity Z-projections of both fluorescence emission channels were produced from the time-lapse acquisitions and single-plane insets were cropped in FIJI. Selected time-frames and insets were further processed with Photoshop CS6 (Adobe).

### Biochemistry

#### Immunoblotting

The protocol used for *total protein extraction* from unfertilised *Drosophila melanogaster* eggs was adapted from Prudêncio and Guilgur, 2015. Eggs from different genetic backgrounds - w^1118^ (negative control), mEGFP-Plk4 (endogenously labelled Plk4), V32-Gal4/ mEGFP-Plk4 (overexpression of labelled Plk4) V32-Gal4/ mGFP-Plk4-ND (positive control, overexpression of non-degradable Plk4, mutated on residues Ser293 and Thr297 (Cunha-Ferreira et al., 2009)) - were collected at 25°C, chemically dechorionated and lysed in ice-cold Lysis Buffer (50 mM Tris-HCl (pH 7.5), 150 mM NaCl, 2 mM EDTA (pH 8), 0.1% Nonidet P-40, 1x Protease inhibitors, 200 mM NaF, 1mM DTT, 150 mM Beta-glycerol phosphate and 1mM Na_3_VO_3_). Samples were snap-frozen in liquid nitrogen and sonicated on ice for 30 sec at about 22% amplitude, using the *Misonix XL 2020 sonicator*. Soluble fractions were cleared by two consecutive rounds of centrifugation at 16000 rcf for 10 min at 4°C, transferring the supernatant into a new Protein LoBind Eppendorf in between, while avoiding the upper lipid layer. The samples were denatured at 99°C for 10 min in 6x SDS Loading Buffer, supplemented with Protease inhibitors. In all cases, 100-120 µg of total protein were run on Invitrogen NuPAGE 4-12% precast Bis-Tris polyacrylamide gels for SDS-Page (Thermo Fisher Scientific) and then transferred onto nitrocellulose membranes for Western Blotting. Membranes were blocked using 5% milk powder in TBS with 0.1% Tween 20 (TBST) for 1h at RT. Primary antibody incubation was performed overnight at 4°C using the following antibodies: anti-GFP rabbit (Abcam, diluted 1:2.000 in 5% milk in TBST) and anti-actin (Sigma, diluted 1:2.000 in 5% milk in TBST). The membranes were washed three times in TBST for 15 min. For detection of GFP-Plk4, secondary antibody anti-rabbit HRP (Bethyl Laboratories) was diluted 1:40.000 (in 2.5% milk in TBST), while for detection of actin, anti-rabbit HRP was diluted 1:20.000 (in 2.5% milk in TBST). The membranes were incubated with secondary antibodies for 1h30min at RT and then washed twice in TBST and once in TBS. For detection of GFP-Plk4, membranes were incubated in WesternBright Sirius HRP substrate (Advansta), while detection of actin was performed by incubating membranes with Pierce ECL Western Blotting Substrate (Thermo Fisher Scientific), following the manufacturer’s instructions. Finally, membranes were exposed for 10-30 s (in the case of GFP-Plk4) and for 3–5 min (in the case of actin) on a Amersham Imager 680 (GE Healthcare) and images were acquired with its CCD camera.

#### mNeonGreen purification

The mNeonGreen coding sequence was cloned with an N-terminus Streptavidin-Binding Peptide (SBP)-Tag and a flexible linker, into the pETMz expression vector (gift from the EMBL Protein Expression & Purification Facility, Heidelberg, Germany), between NcoI and BamHI restriction sites. The 6xHis-Z-tag-TEV-SBP-linker-mNeonGreen protein was expressed in BL21 (Rosetta) Competent *E. coli* at 25°C for 5 hours. The grown liquid culture was harvested and centrifuged at 4000 rpm for 25 minutes, at 4°C. The pellet was ressuspended in ice-cold lysis buffer containing 50 mM K-Hepes (pH 7.5), 250 mM KCl, 1mM MgCl_2_, 1 mM DTT, 7 mM of Imidazole, 1x DNaseI and 1x Protease inhibitors. The sample was applied to a pre-chilled French-press, equilibrated with Lysis buffer, and run twice at a constant pressure (around 12kPa). The cell lysate was collected in a flask on ice and ultracentrifuged at 4°C for 25 min at 50000 rpm using a Ti-70 rotor (Beckman). The protein purification was done through affinity chromatography on a Ni-column (HiTrap chelating HP column 1 ml, GE HealthCare). The column was loaded with a filtered solution of 100 mM nickel chloride, washed extensively with milliQ water and equilibrated with wash buffer (50 mM K-Hepes (pH 7.5), 250 mM KCl, 1mM MgCl_2_, 1 mM DTT, 7 mM of Imidazole). The clarified lysate was applied to the column (at 1.5 ml/min), followed by 200 ml wash buffer. The protein was eluted at 1.5 ml/min with elution buffer: 50 mM K-Hepes (pH 7.5), 250 mM KCl, 1mM MgCl_2_, 1 mM DTT, 400 mM of Imidazole. 1 ml sample fractions were collected and kept at 4°C. The most concentrated samples were pooled together and their N-terminus 6xHis-Z-tag was cleaved with TEV protease overnight at 4°C by treating with 150U TEV/mg of protein. The following day, the cleaved protein was passed through a column for size-exclusion chromatography to remove contaminants, the cleaved tag and the TEV protease (with Tiago Bandeiras at IBET, Oeiras, Portugal). Additionally, the elution buffer was exchanged to a storage buffer: 50 mM K-Hepes (pH 7.8), 100 mM KCl, 2 mM MgCl_2_, 1 mM DTT, 1 mM EGTA. The HiLoad Superdex 75 16/60 (GE HealthCare) gel filtration column was equilibrated with storage buffer for 1hour. The sample was spun at 15000 rpm for 15 min at 4°C and the clear fraction was applied to the gel filtration column coupled to an AKTA device at 1 ml/min. The cleaved mNeonGreen protein was concentrated approximately 5 times using Amicon 10K Centrifugal filters. Pure glycerol was added at 5% v/v and small aliquots were snap-frozen in liquid nitrogen and stored at −80°C.

#### Plk4 titration in cytoplasmic extract

Plk4 dilution was accomplished by mixing cytoplasm from flies with different genetic composition. Unfertilised eggs collected from females overexpressing Plk4 in the germline (genotype: V32-Gal4/ pUb-Spd2–GFP; Jupiter-mCherry/pUASp-GFP–Plk4) were homogenised in unfertilised eggs from females without the transgenic pUASp element (genotype: V32-Gal4/ pUb-Spd2–GFP; Jupiter–mCherry), where all components are at wild-type levels, specifically diluting overall Plk4 concentration in the cytoplasm. Different final Plk4 concentrations were achieved by mixing Plk4 overexpression:wildtype eggs at the following ratios: 6:0 (“1” relative Plk4 concentration, control); 3:3 (“0.5” relative Plk4 concentration); 2:4 (“0.33” relative Plk4 concentration) and 1:5 (“0.16” relative Plk4 concentration). Small explants were produced from the cytoplasmic mixtures and images were acquired for 40 minutes. All time-lapse acquisitions within this section were performed at 1 minute time-interval with 0.45 µm optical sections, using a Plan Apo VC 60x 1.2 NA water objective.

### Fluorescence Correlation Spectroscopy (FCS) data acquisition and analysis

#### Standard rhodamine 6G calibration

All FCS measurements were performed on a point-scanning confocal microscope (Zeiss LSM780 Confocor3) equipped with a UV-VIS-IR C Achromat 40X 1.2 NA water-immersion objective and a gallium arsenide detector array wavelength selected between 491-561nm. Before each experiment the system was aligned using a high concentration and calibrated using a low concentration Rhodamin 6G solution in water. The known diffusion coefficient of rhodamine 6G (410 µm^2^/s) (Majer and Zick, 2015) allowed us to determine the lateral beam waist (w_xy_ = 232 nm) and the structure factor (S = 5.77) of the focused laser (Point Spread Function, PSF). The resultant volume of illumination is calculated through:

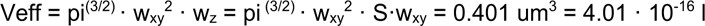

The values for w_xy_ and S were used as constants in the subsequent model-based fittings of the autocorrelation functions (ACF) and the volume was used to calculate the concentration (see below).

#### Calibration with purified mNeonGreen

mNeonGreen fluorescent tag was first measured in a cytoplasm-compatible buffer. Fluorescence intensity in time (I(t)) was recorded as 6 iterations of 10s. Each 10s trace was autocorrelated into an ACF, G(τ), using the Zeiss onboard autocorrelator which calculates the self-similarity through:

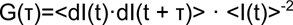

Here <> denotes the time-average, dI(t)=I(t)-<I(t)> and τ is called the timelag. The resulting G(τ) curves of the fluorophores in buffer were readily fitted using a regular 3D diffusion model:

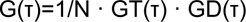

where N reflects the number of moving particles in the confocal volume and GT(τ) is the correlation function associated to blinking/triplet kinetics:

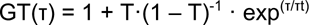

Where T is the fraction of molecules in the dark state and τt the lifetime of the darkstate. GD(τ) is the correlation function associated to diffusion which in this case is simple Brownian diffusion in 3D:

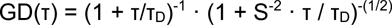

These fittings allowed us to measure the number of molecules in the confocal volume and therefore their brightness (<I(t)> / N) together with the characteristic diffusion times (τ_D_).

The above model fit is based on the assumption that there are only two characteristic timescales generating the ACF. In order to get a model free estimate of the number of timescales involved we used a Maximum Entropy Method based fitting (MEMfit) of the combined and normalised ACFs of each experiment. MEMfit analyses the FCS autocorrelation data in terms of a quasicontinuous distribution of diffusing components making it an ideal model to examine the ACF of a highly heterogeneous system without prior knowledge of the amount of diffusing species.

To be able to quantify the brightness of individual fluorescent tags in an embryo the purified mNeonGreen was injected into pUb-RFP-**β**2-Tubulin dechorionated embryos. An anomalous coefficient had to be included to fit the resultant ACF:

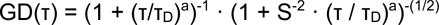

For simple Brownian diffusion a = 1 and the fit function is identical to the one used to fit the fluorophores in buffer. However, for fluorophores injected into the cytosol of embryos the fitting algorithm gave an anomalous coefficient of a = 0.8. An anomalous coefficient smaller than 1 indicates constrained diffusion and could be caused by the more crowded environment in the yolk. In addition, the large amount of (uncorrelated) autofluorescence generated by the yolk leads to an underestimation of the brightness therefore requiring a background correction factor. The background values were determined per excitation power from embryos lacking the Plk4 reporter. If the background itself does not autocorrelate it has no influence on the obtained timescales in the data. Nevertheless, the background will impact the absolute number, N, and consequently also the calculated brightness. Therefore, all the measurements were background corrected via:

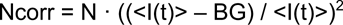

Where BG is the measured background from embryos lacking the reporter fluorophore. Consequently the corrected brightness was calculated as:

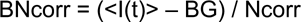

Finally, any 1 millisecond-binned intensity trace that contained changes in average intensity (most likely arising from yolk spheres moving through the confocal spot during the measurement) were discarded from further analysis.

#### mNeonGreen-Plk4 measurements in embryos

For the measurements of mNeonGreen-Plk4, embryo staging was done based on the pUb-RFP-**β**2-Tubulin reporter. We chose embryos at blastoderm stage, in division cycles 10 or 11. Before each FCS acquisition series, a large field-of-view image of the embryo was acquired. Six different, 10 seconds long intensity traces were measured at the inter-nuclear cytoplasmic space of the syncytium. The 10s measurement was long enough to obtain sufficient passage events and short enough to avoid each trace to be contaminated by events that do not arise from mNeonGreen-Plk4 diffusing in the cytosol.

From these measurements, the MEMfit method on the normalised ACF indicates three timescales for the tagged-Plk4 molecules. A first timescale of 5-50 µs corresponding to the triplet state dynamics that were similarly found in both the buffer as well as from fluorophores injected in the embryo. A second timescale of about 0.8ms, most likely coming from the diffusion of a Plk4 monomer (see similarity to mNeonGreen monomer in cytosol). And a third timescale of diffusion that is much slower, 9ms. In order to fit the ACFs the diffusional part of the fit function was associated with two components:

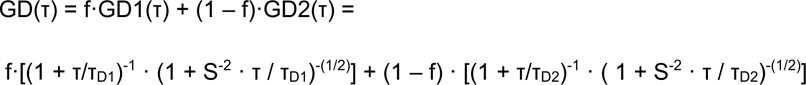

The fraction f corresponds to the fast diffusing Plk4. The Diffusion Coefficient of each of the components can be calculated from the diffusion timescales τD via:

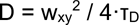

### In vitro experiments

#### *Drosophila melanogaster* cell culture

Drosophila (DMEL) cells were cultured in Express5 SFM (GIBCO, USA) supplemented with 1x L-Glutamine-Penicillin-Streptomycin. Double-stranded RNA (dsRNA) synthesis was performed as previously described (Bettencourt-Dias et al., 2004). DMEL cells were plated and treated for 12 days with 40 μg dsRNA against Plk4 or mCherry (control), replacing the dsRNA every 4 days. Cells were fixed at day 10 to confirm centriole depletion and treatment with dsRNA agains PCM was initiated. Cells were then treated for 6 days with different amounts and combinations of dsRNA: 80 μg mCherry alone, 20 μg of individual PCM components – Cnn, Asl, D-Plp, Spd2 or γ-tubulin 23C – or combinations of two – Cnn + Spd2,Cnn + D-Plp or Spd2 + D-Plp – three – Cnn + Spd2 + D-Plp - or four of these molecules – Cnn + Asl + D-Plp + Spd2 (referred to as ‘All PCM’). Primers used for dsRNA synthesis are listed in Suppl. Table S5.

#### Immunostaning and imaging of *D. melanogaster* cultured cells

DMEL cells were plated onto clean glass coverslips and allowed to adhere for 1 hour and 30 min. The media was removed and cells were fixed at −20°C for 10 min in chilled methanol. Cells were permeabilised and washed in D-PBSTB (1x Dulbecco’s Phosphate Buffered Saline pH 7.3, with 0.1% Triton X-100 and 1% BSA) for 1 hour. Cells were incubated overnight at 4°C with primary antibodies – rat anti-Sas4 (dilution 1:500) kindly provided by David Glover (University of Cambridge, UK) and rabbit anti-CP110 (dilution 1:10000; Metabion) – diluted in D-PBSTB. Cells were washed in D-PBSTB and incubated for 1hour 30 min at room temperature with secondary antibodies – donkey anti-rat Alexa 555 (dilution 1:1000; Molecular Probes) and donkey anti-rabbit Alexa 647 (dilution 1:1000; Jackson Immunoresearch Laboratories) – and DAPI (dilution 1:200) in D-PBSTB. Cells were washed and mounted with Dako Faramount Aqueous Mounting Medium (S3025, Agilent).

Cell imaging was conducted on a Nikon Eclipse Ti-E microscope equipped with a Yokogawa CSU-X1 Spinning Disk confocal scanner. Images were recorded with a EMCCD Photometrics 512 camera. Optical sections of 0.3 µm thickness were acquired with a Plan Apo 100x 1.49 NA oil immersion objective using a piezoelectric stage (737.2SL, Physik Instrumente), controlled by Metamorph 7.5 software. Centriole number was scored in 300 cells per treatment, per independent experiment. Data is presented as average (with standard error mean, S.E.M.) of two independent experiments. We tested all counts with a Chi-square test against the null-hypothesis that the outcome is random. Then, each 16d test condition was compared to the 16d mCherry control condition with a 2-proportions Z-test and H0 that the proportions of cells with centrioles are equal versus HA that the proportion in the test is smaller. The significance level for multiple testing was Bonferroni corrected. Significance level was p = 0.01. All images were processed with ImageJ (NIH, USA) and Adobe Photoshop CS6 (Adobe Systems, USA), and the final figures were assembled in Adobe Illustrator CS6 (Adobe Systems, USA).

## Acknowledgments

We would like to thank the Central Imaging and Flow Cytometry Facility (CIFF) at the National Centre for Biological Sciences (NCBS) in Bangalore, where all FCS experiments were performed.

We acknowledge the technical support of IGC’s Advanced Imaging Facility (AIF), in particular Gabriel Martins and Nuno Pimpão Martins. IGC’s AIF is supported by the national Portuguese funding ref# PPBI-POCI-01-0145-FEDER-022122, co-financed by Lisboa Regional Operational Programme (Lisboa 2020), under the Portugal 2020 Agreement, through the European Regional Development Fund (FEDER) and Fundação para a Ciência e a Tecnologia (FCT, Portugal).

We thank IGC’s Fly Facility and Fly Transgenesis Facility supported by Congento Congento LISBOA-01-0145-FEDER-022170, co-financed by FCT (Portugal) and Lisboa 2020, under the PORTUGAL 2020 agreement (European Regional Development Fund). Transgenic fly stocks were obtained from Bloomington Drosophila Stock Center (NIH742 P40OD018537).

We acknowledge financial support from Boehringer Ingelheim Fonds PhD Fellowship awarded to C. Nabais, Human Frontiers Science Program (HFSP) Young Investigator Grant (RGY0083/2016) awarded to I.A. Telley supporting J. de–Carvalho, the Fundação para Ciência e a Tecnologia (FCT) supporting I.A. Telley (Investigador FCT IF/00082/2013), the EU FP7-PEOPLE-2013-CIG (N° 818743) awarded to I.A. Telley, ERC-2010-StG-261344-CentriolStructure&Number and an ERC-2015-CoG-683258-Birth&Death awarded to M. Bettencourt-Dias, and the Gulbenkian Foundation (FCG). T.S. van Zanten acknowledges an EMBO fellowship (ALTF 1519-2013) and a NCBS Campus fellowship. S. Mayor acknowledges a JC Bose Fellowship from DST (Government of India), support from the NCBS-Max Planck Lipid Centre, a grant from HFSP RGP0027/2012, and support from Wellcome Trust/DBT India Alliance Margdarshi Fellowship (IA/M/15/1/502018).

We thank Tiago Bandeiras at Instituto de Biologia Experimental e Tecnológica (IBET), Oeiras, for the gel filtration chromatography conducted in his facility with Micael Freitas.

We thank Tomer Avidor-Reiss, Daniel St Johnston, Yoshihiro Inoue and Jordan Raff for sharing transgenic fly lines. We thank David Glover, Jordan Raff and Tim Megraw for providing antibodies.

We thank members of the Cell Cycle Regulation lab at IGC for giving feedback to earlier versions of the manuscript.

Finally, we wish to thank the anonymous reviewers and the editor for helpful comments which greatly improved the manuscript.

## Author contributions

Conceptualization: CN, IAT, MBD

Methodology: CN, JdC, IAT (egg explant assay); CN, TvZ (FCS measurements)

Software: DP, JC (design and implementation of model simulations)

Validation: CN, JdC, DP, TvZ, SM, JC, IAT, MBD

Investigation: CN, JdC (performing data collection egg explants); CN, TvZ (performing data collection in FCS measurements); DP, JC (collecting in silico data)

Analysis: CN, IAT (experimental data from egg explants, eggs and cell culture); TvZ, SM (FCS data); DP, JC (theoretical model)

Resources: CN (CRISPR fly line, vectors and plasmid design, recombinant protein purification); PD (genotyping); IAT (design of micromanipulation microscope)

Visualization of data: CN, DP, TvZ, IAT

Writing – original draft: CN

Writing – review and editing: CN, SM, JC, IAT, MBD

Supervision and coordination: IAT, MBD

---

**Supplementary Figure 1 (In support of Figures 1 & 2).**
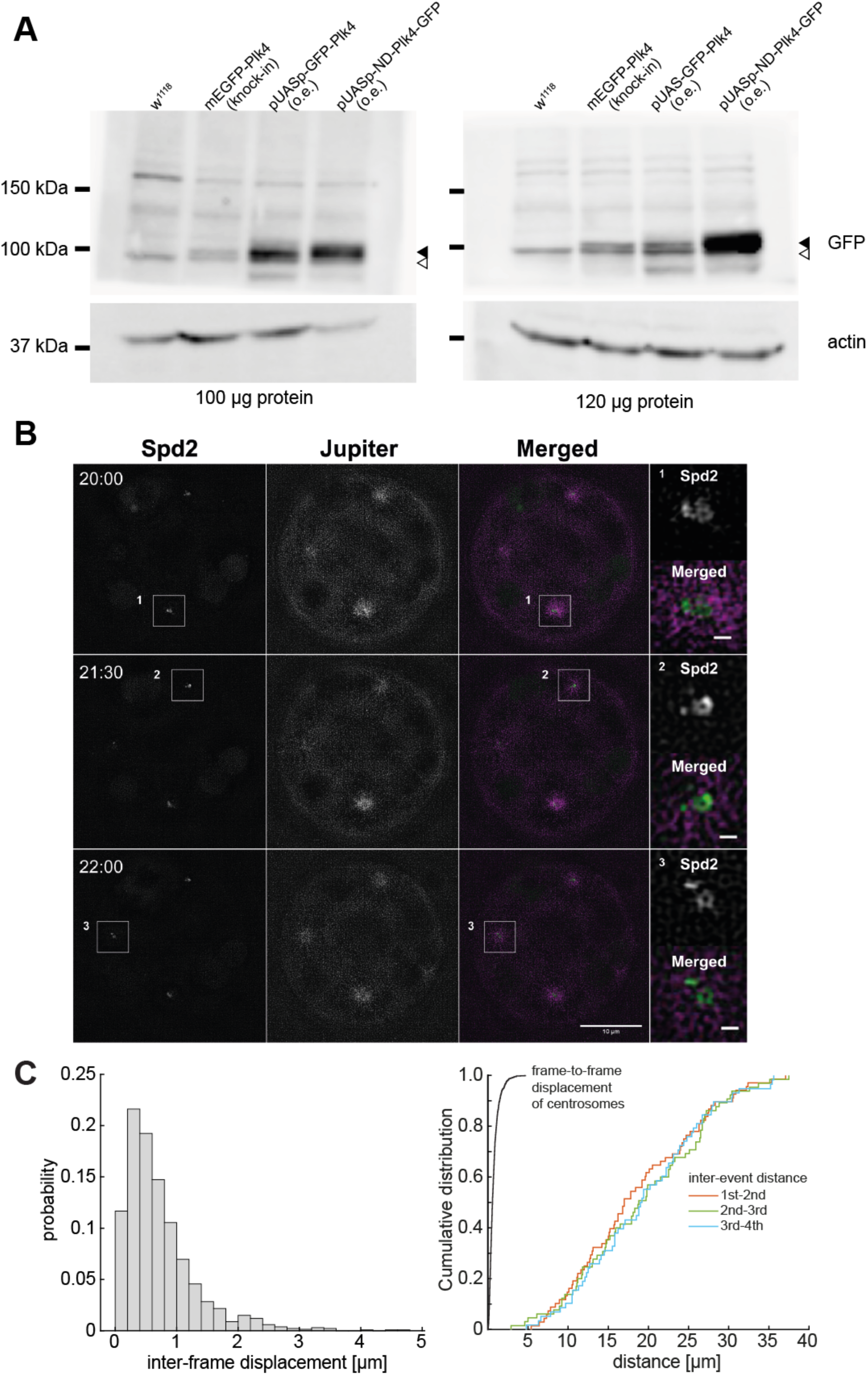
**(A)** Western blot analysis of Plk4 concentration for endogenous expression and for overexpression constructs. We emphasise that the detection of endogenous Plk4 with a Western blot approach is extremely challenging. In fact, most studies so far have only detected Plk4 by means of affinity-tag or fluorescent reporter and/or under an overexpression scenario. We were able to visualise the endogenous Plk4 tagged with an mEGFP tag (CRISPR fly line, generated in the same way as the neon-green PLK4 represented in Suppl. Fig. 4A,B). PLK4 overexpression was visualised with a GFP–PLK4 overexpression (o.e.) construct, whose extract shows similar centriole biogenesis results as the PLK4 non tagged construct used in most of this manuscript (now shown). We also made extract from flies overexpressing non-degradable PLK4 which accumulates in embryos and serves as a positive control (pUASp-ND-PLK4–EGFP, Cunha-Ferreira et al., 2013). WT embryos were loaded as negative controls. Black arrowhead points to GFP-tagged Plk4 constructs, while white arrowhead points to an unspecific signal also present in WT embryos. We register 3.2±1.9 times higher Plk4 concentration in the extract of embryos overexpressing Plk4 as compared to the wildtype (from four experiments). Our conservative interpretation is that overexpression is less than a magnitude of the endogenous concentration. This result is in line with our dilution results, where at 1/5 dilution of the Plk4-overexpressing extract, we see very few de novo events (Fig. 4). **(B)** Visualisation of centrosome biogenesis in a *Drosophila* egg extract by 3D-Structured Illumination Microscopy (3D-SIM). Maximum-intensity Z-projections from a time-lapse acquisition of an unfertilised egg explant overexpressing Plk4. Centrioles (insets) are detected as barrel-shaped structures surrounded by the PCM component Spd2 (green) associated with a microtubule array (magenta), reported by the microtubule associated protein Jupiter. Insets are single-plane images of three different centrosomes. Scale-bar, 0.5 μm. Centrioles formed de novo can duplicate. Time is reported as min:sec. **(C)** Comparison of centrosome movement versus distance between registered biogenesis. **Left**: histogram of frame-to-frame (instantaneous) displacements of 1^st^ event centrosomes. Most of the centrosomes performed random movement and only in rare cases they moved away in a directed fashion from the explant boundaries. **Right**: Cumulative distribution functions of frame-to-frame displacement (black) in comparison with all subsequent inter-event distances as presented in Fig. 3. This comparison shows that the probability for a biogenesis event to quickly displace a distance typically seen between biogenesis events is extremely low. Any subsequent event after the first biogenesis is unlikely a “duplication-and-run” event.

**Supplementary Figure 2 (In support of Figure 3).**
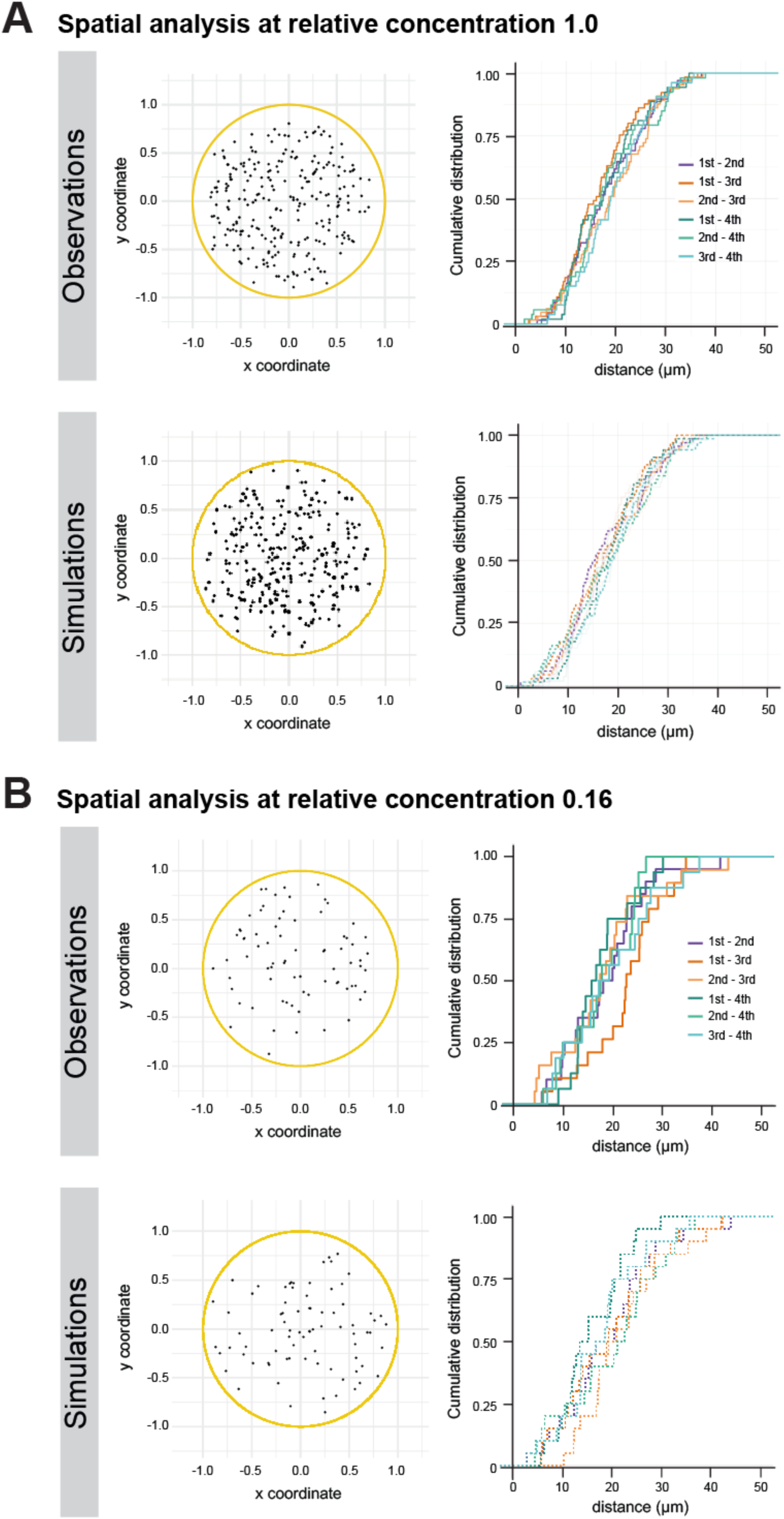
**(A)** Spatial analysis of de novo centriole biogenesis in fly explants. **Left:** 2D Z-projections of the positions of centrioles at the moment they were first detected in the explants - 254 centrioles measured in 68 explants (“Observations”) and 272 centrioles from 68 *in silico* explants (“Simulations”). All coordinates were normalized to the measured explant diameter. **Right:** Distributions of observed and simulated inter-event distances measured in 3D for the first four centrosomes formed de novo in the explants. **(B)** Spatial analysis of de novo centriole biogenesis at lower Plk4 concentration. **Left:** Z-projections of the positions of centrioles at the moment they were first detected in the explants – 75 centrioles measured in 20 explants (“Observations”) and 80 centrioles from 20 *in silico* explants (“Simulations”). All coordinates were normalized to the measured explant diameter. **Right:** Distributions of observed and simulated inter-event distances measured in 3D for the first four centrosomes formed de novo in the explants, at the lowest Plk4 overexpression (“0.16” relative concentration of Plk4). The grey envelope indicates the 95% Confidence Interval (from quantile 0.025 to 0.975) for the simulated data.

**Supplementary Figure 3 (In support of Figure 4).**
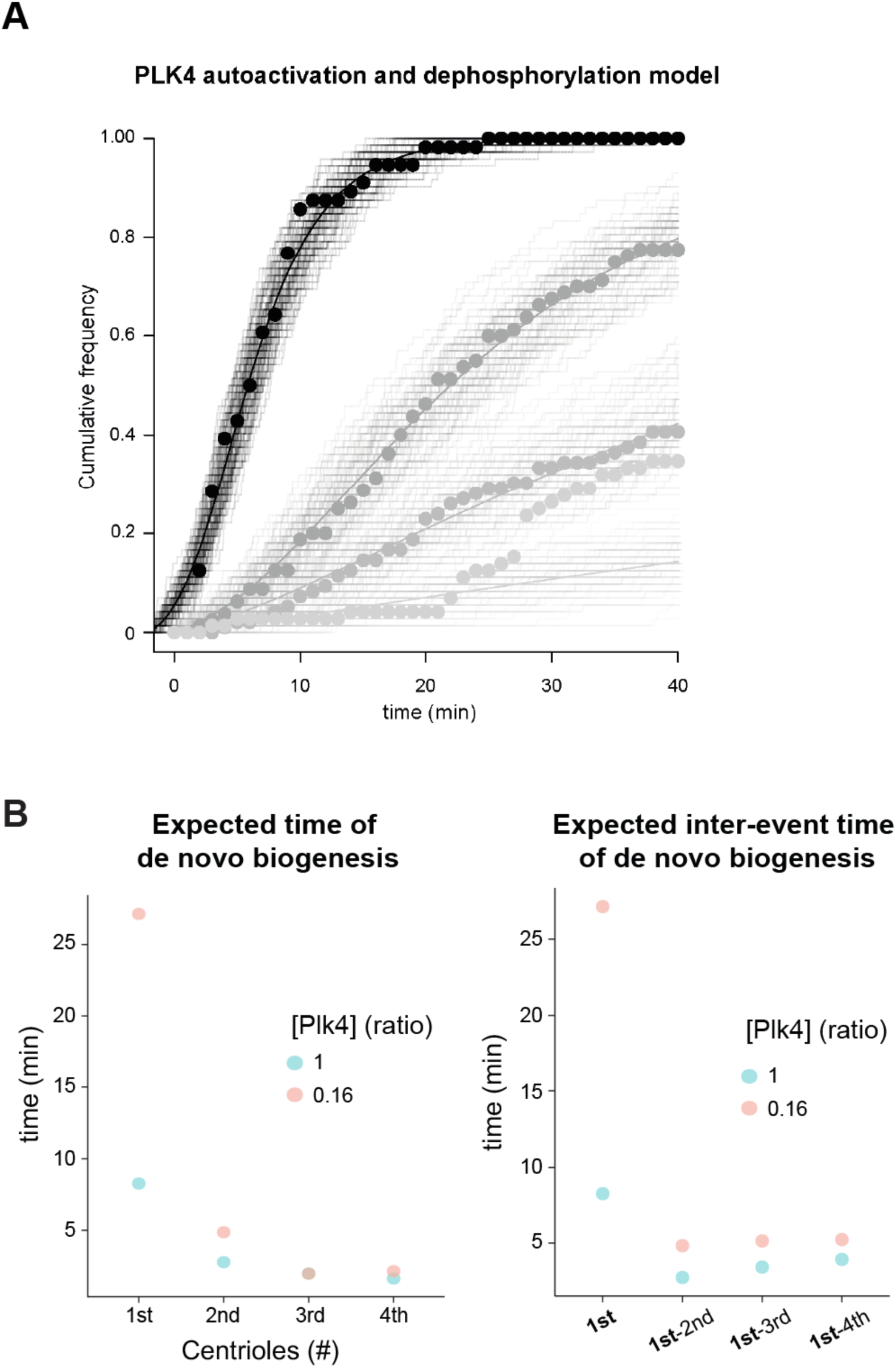
**(A)** Fitting of Plk4 autoactivation and dephosphorylation model to data measured in explants at different Plk4 concentrations. The grey gradient represents different concentrations of Plk4. The different concentrations were prepared experimentally by mixing the cytoplasm from high overexpression eggs (taken as the unit “1”, black) with cytoplasm from wild-type eggs, in different proportions such that the dilutions are “0.5”, “0.33” and “0.16” relative concentrations. The dots are the relative frequency of explants containing at least one de novo formed centriole for the different concentrations of Plk4: “1” (N = 56), “0.5” (N = 62), “0.33” (N = 39) and “0.16” (N = 25). The lines are the solution of the model of Plk4 trans-autophosphorylation (Fig. 4A). The continuous lines are the solution of the ordinary differential equation model and the staircase lines are the results of stochastic simulations under the same parameter settings. The Plk4 activity in the higher concentration (denoted K) was adjusted, whereas the activities in the dilutions were set in relative terms (0.16K, 0.33K and 0.5K). The modelling and simulations, as well as the remaining parameters and values are described in section the Methods (Statistics and mathematical modelling). Notice that as Plk4 concentration decreases, so does the number of droplets where centriole biogenesis occurs within 40 minutes of time-lapse recording. **(B)** Temporal kinetics of de novo centriole biogenesis at different concentrations of Plk4 investigated in Figure 4. **Left:** Estimation of the mean centriole biogenesis times at the highest Plk4 concentration (“1”, in blue) and at the lowest Plk4 overexpression (“0.16”, in orange) by ML estimation (MLE) fitting of a simple exponential model. **Right:** Estimation of the waiting time until the first de novo event and inter-event time between the first and subsequent de novo events, at high (“1”, in blue) and the lowest (“0.16”, in orange) concentration of Plk4.

**Supplementary Figure 4 (In support of Figure 5).**
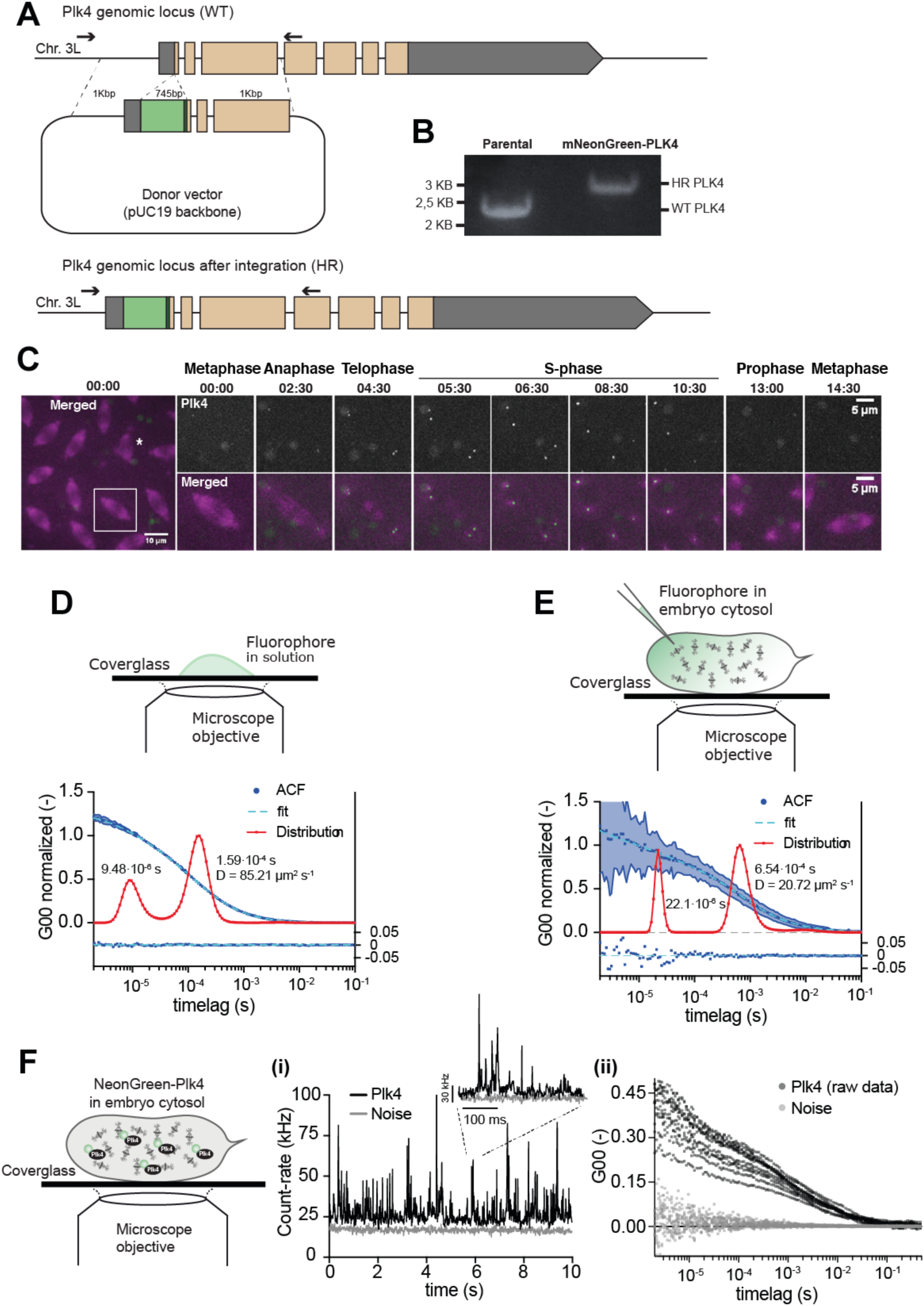
**(A)** Insertion of a fluorescent tag into *Drosophila* Plk4 endogenous locus. Schematic representation of the wildtype dmPlk4 locus (WT) and of the dmPlk4 locus after successful tag integration (HR). A donor plasmid carrying the mNeonGreen reporter and a small linker (dark green) flanked by 1 Kbp homology arms was used for homologous recombination. The UTRs are shown in grey and the coding sequences are depicted in orange. The arrows indicate the position of the screening primers dmPLK4 5UTR 3 FW and dmPLK4 1exon Rev, which are located outside the homology arms. The same strategy was used for neongreen and GFP tags, generating two lines that were used at different parts of this manuscript. **(B)** Integration of a fluorescent tag into Plk4 endogenous locus (HR Plk4) causes a migration shift of the PCR product in the agarose gel compared to the untagged Plk4 locus (WT Plk4). **(C)** Maximum intensity Z-projections from a time-lapse video of a syncytial *D. melanogaster* embryo expressing endogenous mNeonGreen-Plk4 (green) and microtubule reporter RFP-**β**-tubulin (magenta). Plk4 localises at the centrosomes (high intensity tubulin spots) in interphase. Larger green dots result from yolk auto-fluorescence. At timepoint t=00:00 the embryo is in metaphase of nuclear cycle 11. The insets show the progression of a single nucleus and its daughters, throughout one cell-cycle. The cell-cycle stage is indicated above each image. Time is reported as min:sec. The asterisk indicates an abnormal mitotic spindle. **(D)** FCS measurements of purified mNeonGreen fluorophore in a buffer supporting viability of the cytoplasm (Telley et al 2013). **(E)** FCS measurements of mNeonGreen after injection into the cytosol of syncytial embryos expressing RFP-Tubulin. The graphs show the normalised, fitted Autocorrelation Functions (ACF, blue dots and light-blue curve), with standard deviation (shaded area) and Maximum Entropy Method (MEM) Fit (red line). The time lags (diffusion times) determined using the two fitting methods shown next to the MEM-fit curves are in agreement. The peak at the fast timescale corresponds to the triplet state of the fluorophore (9.48×10^−6^ s in solution; 22×10^−6^ s in the cytoplasm), whereas the second peak in the slower timescale corresponds to the 3D diffusion of mNeonGreen, from which a diffusion coefficient D was calculated (1.59×10^−4^ s, D = 85.21 μm^2^/s in solution; 6.54×10^−4^ s, D = 20.72 μm^2^/s in the cytoplasm). The residuals obtained from the best fit are shown below the ACF graphs. **(F)** Single-molecule mNeonGreen–Plk4 quantifications in the cytosol of the syncytial fly embryo. **(i)** Intensity traces of mNeonGreen– Plk4 (black) and background noise (grey). Of note, intensity bursts of mNeonGreen–Plk4 are well distinguishable from background noise (inset). **(ii)** Raw auto-correlation functions (ACF) from multiple independent FCS measurements. While the intensity of background acquisitions as measured in RFP–Tubulin expressing embryos does not auto-correlate, traces from mNeonGreen–Plk4 expressing embryos exhibit significant autocorrelation.

**Supplementary Figure 5 (In support of Figure 6).**
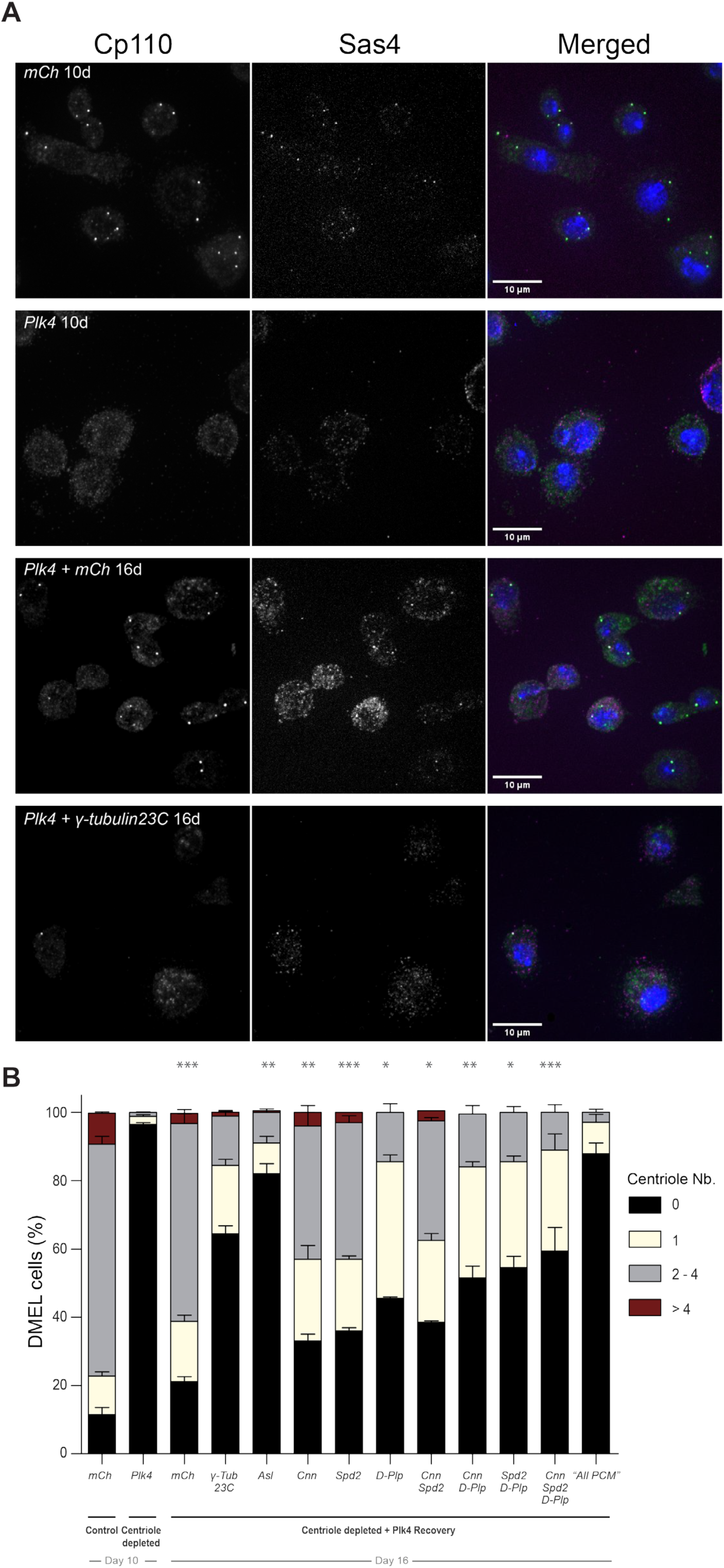
Centriole de novo biogenesis is partially impaired in PCM-depleted DMEL cells. Maximum intensity Z-projections of DMEL cells treated with RNAi against Plk4 or mCherry (mCh) for 10 days. Cells treated with RNAi against Plk4 gradually lose centrioles during proliferation. After 10 days, centriole-depleted cells were allowed to recover Plk4 translation while simultaneously treated for four days with RNAi against individual PCM components – Cnn, Asl, D-Plp, Spd2 or γ-tubulin 23C – or combinations of these molecules previously shown to be essential for PCM assembly in cycling cells - Cnn + Spd2, Cnn + D-Plp, Spd2 + D-Plp or Cnn + Spd2 + D-Plp (Gomez-Ferreria et al., 2007; Conduit et al., 2014; Lerit et al., 2015; Feng et al., 2017; Citron et al., 2018). Additionally, we depleted all four PCM components – Cnn + Asl + D-Plp + Spd2 (referred to as “All PCM”) - required for PCM maintenance (Pimenta-Marques et al., 2016). The panels show centriole-depleted cells treated with RNAi against mCherry (recovering centriole normal number) and γ-tubulin 23C (abnormal centriole number). Cells were stained with centriolar markers Sas4 (magenta) and Cp110 (green); and DAPI-stained (DNA, in blue). Note that it is very common for a small fraction of untreated DMEL cultured cells (“wildtype”) to have either too many (more than 4) or too few (less than 2) centrioles (Bettencourt-Dias et al., 2005). This is found in most cell lines from *Drosophila melanogaster* as they are permissive to those changes, as in contrast to vertebrate cells, there is no p53-dependent cell-cycle arrest in the presence of numerical centrosome abnormalities. **(B)** Quantification of centriole number per cell after 10 and 16 days of RNAi treatment. Data are the average of two independent experiments (with standard error of the mean - S.E.M.). Superscripts ‘*’ denote statistical significance in treatments, where *, ** and *** indicate p< 0.05, 0.01, 0.001 (Pearson’s χ^2^ test and 2-proportions Z-test).

**Supplementary Figure 6 (In support of Figure 6).**
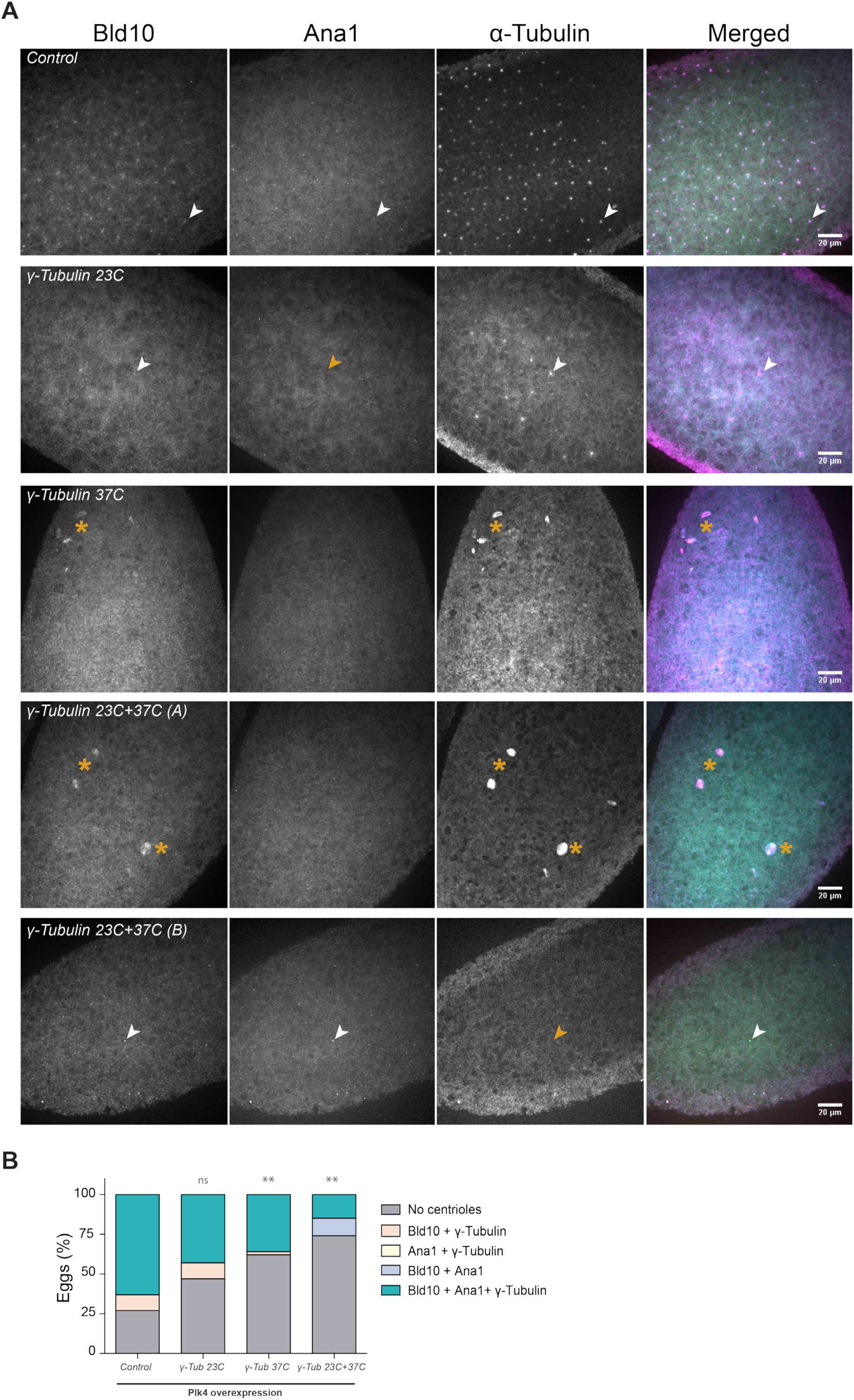
De novo centriole biogenesis is partially impaired in unfertilised eggs overexpressing Plk4 and depleted for Gamma-tubulin. **(A)** Maximum intensity Z-projections of unfertilised eggs overexpressing Plk4 alone (Control) or together with RNAi against γ-tubulin 23C, γ-tubulin 37C or both. Eggs were stained with Bld10 (cyan), Ana1 (yellow) and tyrosinated α-tubulin (magenta). Centrioles (arrowheads) were identified by co-localisation of at least two of these markers. Yellow arrowheads depict centrioles for which one of the centrosomal proteins is not detected. Yellow asterisks reveal putative meiotic defects, previously described to occur in oocytes from γ-tubulin 37C mutant females (Tavosanis et al., 1997). **(B)** Quantification of unfertilised eggs with de novo centriole assembly driven by Plk4 overexpression and detected by the combination of two or three of either Bld10, Ana1 or tyrosinated α-tubulin. ** and *** indicate p< 0.01 and 0.001 respectively, Pearson’s χ^2^ test and 2-proportions Z-test.

## SUPPLEMENTARY TABLES

**Supplementary Table 1.**
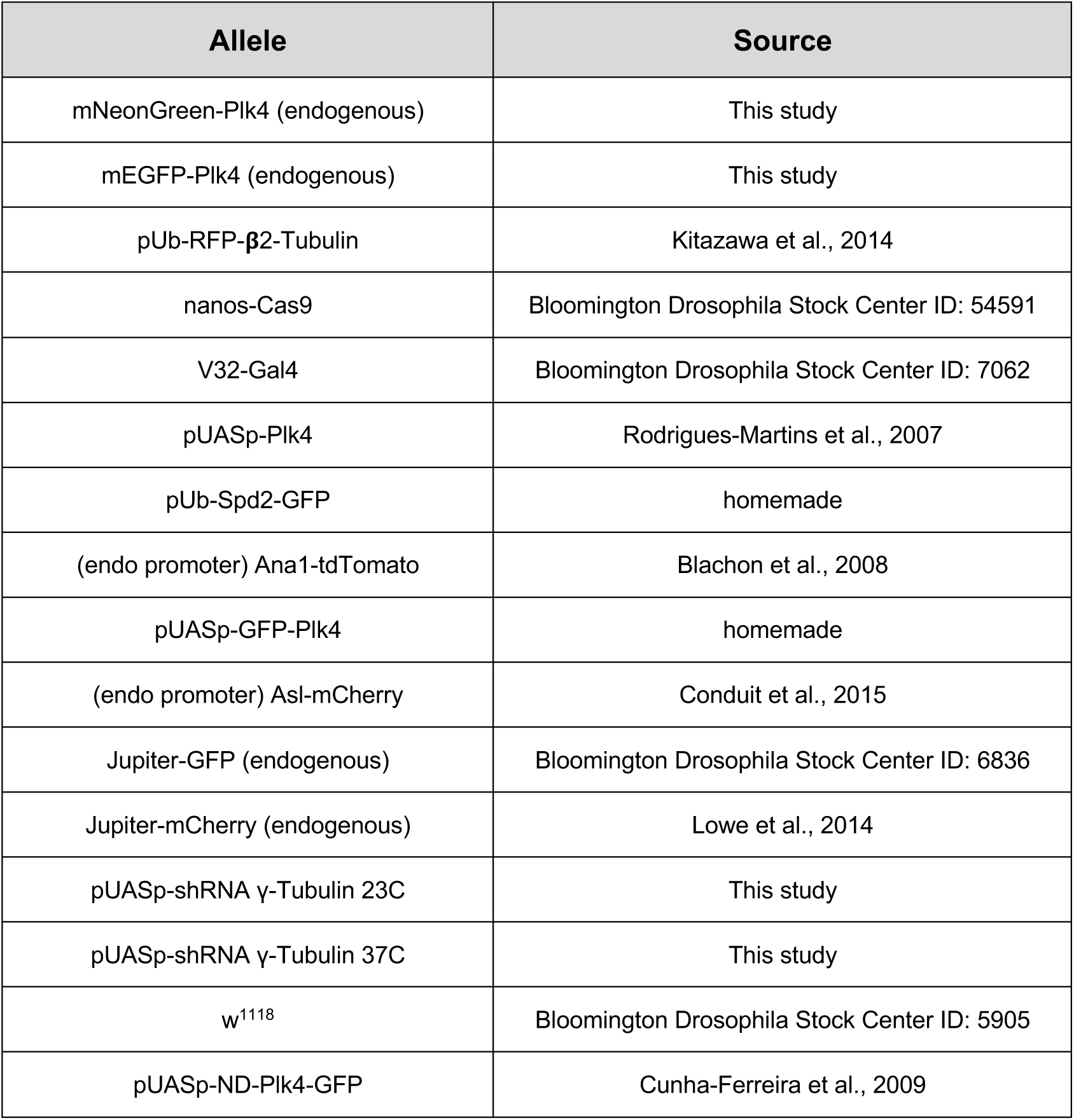
*D. melanogaster* strains generated and/or used in this study.

**Supplementary Table 2.**
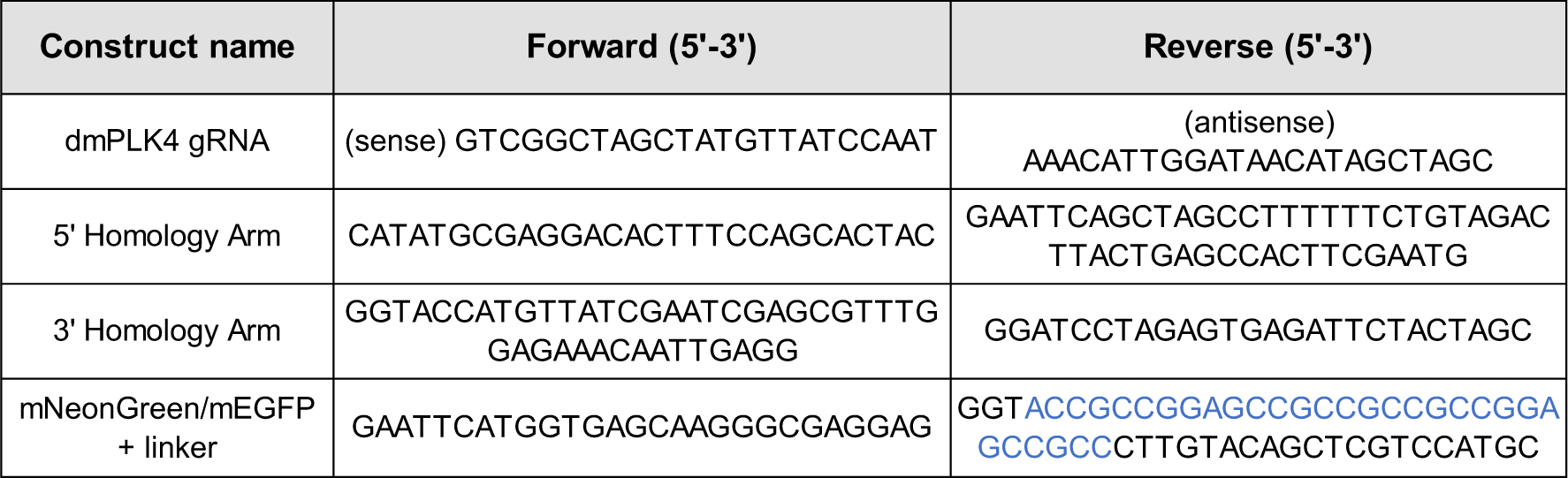
List of oligonucleotides used for CRISPR-mediated knock-in of mNeonGreen and mEGFP into the endogenous *Drosophila melanogaster* Plk4 locus. The guide RNA (gRNA) was used to the target the genome editing at dmPlk4 N-terminus. The three combinations of primers were used to clone the donor vector with the mNeonGreen fluorescent reporter. A short flexible linker (highlighted in blue) was placed between the coding sequences of mNeonGreen and Plk4.

**Supplementary Table 3.**
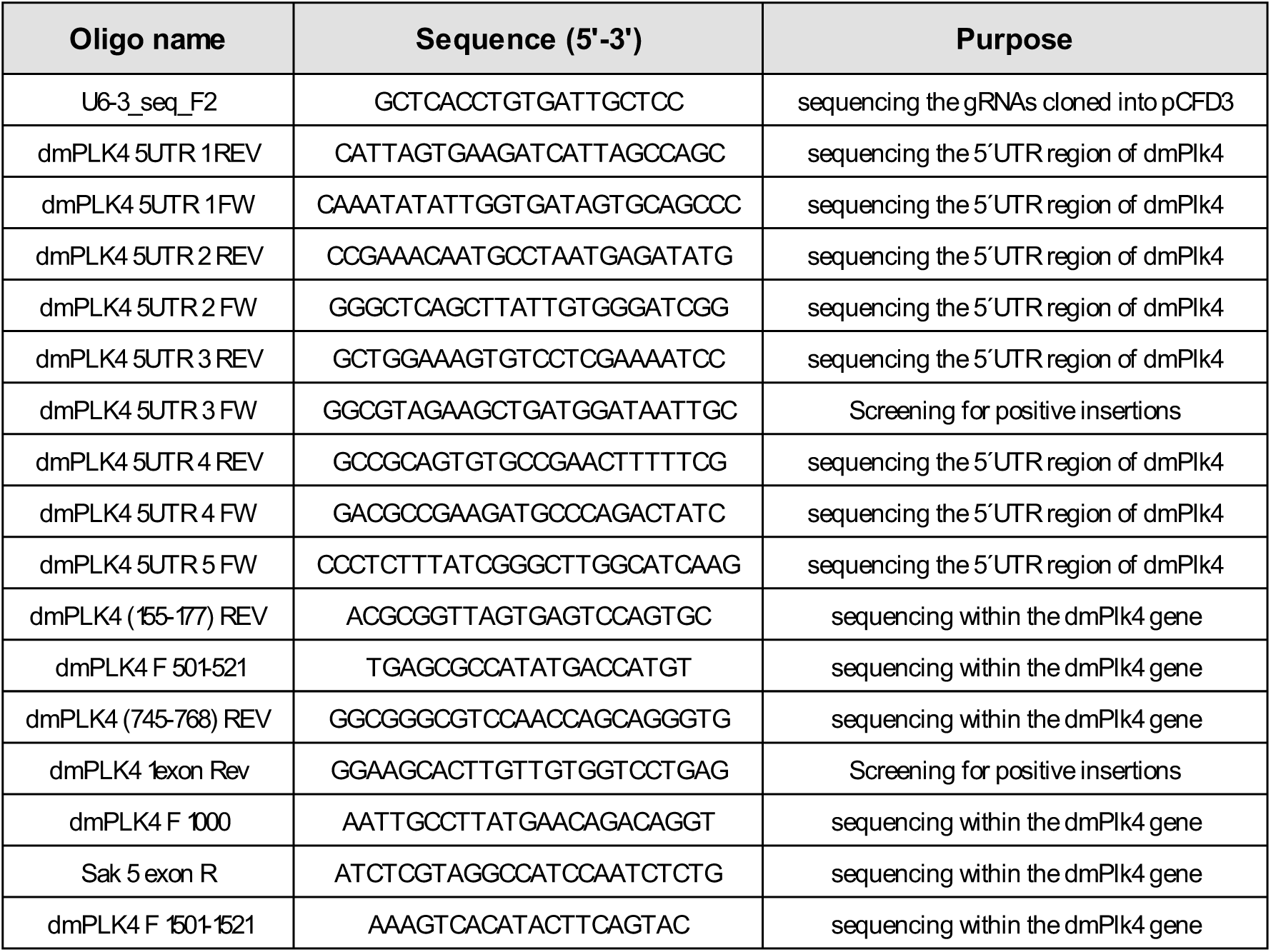
Sequencing and screening primers used to check the mNeonGreen-Plk4 and mGFP-Plk4 lines generated in this study.

**Supplementary Table 4.** FCS Parameters determined from the model-based fittings. Total number of measurements and embryos analysed and diffusion model applied to each experimental condition. According to the model, either one or two diffusion components were determined and their characteristic timescales and diffusion coefficients calculated. The fraction of each diffusing pool is presented as a percentage.

| | No. of measurements/<br>No. of embryos | Diffusion model | timescale 1<br>(ms) | Diffusion coeff 1<br>($\mu\text{m}^2/\text{s}$) | Fraction of $\tau\text{D1}$ (%) | timescale 2<br>(ms) | Diffusion coeff 2<br>($\mu\text{m}^2/\text{s}$) | Fraction of $\tau\text{D2}$ (%) |
| --- | --- | --- | --- | --- | --- | --- | --- | --- |
| mNeonGreen in solution | 24 | 1 component 3D | 0.15 | 85.2 | 100 |  |  |  |
| mNeonGreen in the cytosol | 85 / 9 | 1 component anomalous 3D | 0.65 | 20.7 | 100 |  |  |  |
| mNeonGreen-Plk4 in the cytosol | 147 / 11 | 2 component 3D | 0.79 | 17.2 | 52.3 | 9.11 | 1.49 | 47.7 |

**Supplementary Table 5.**
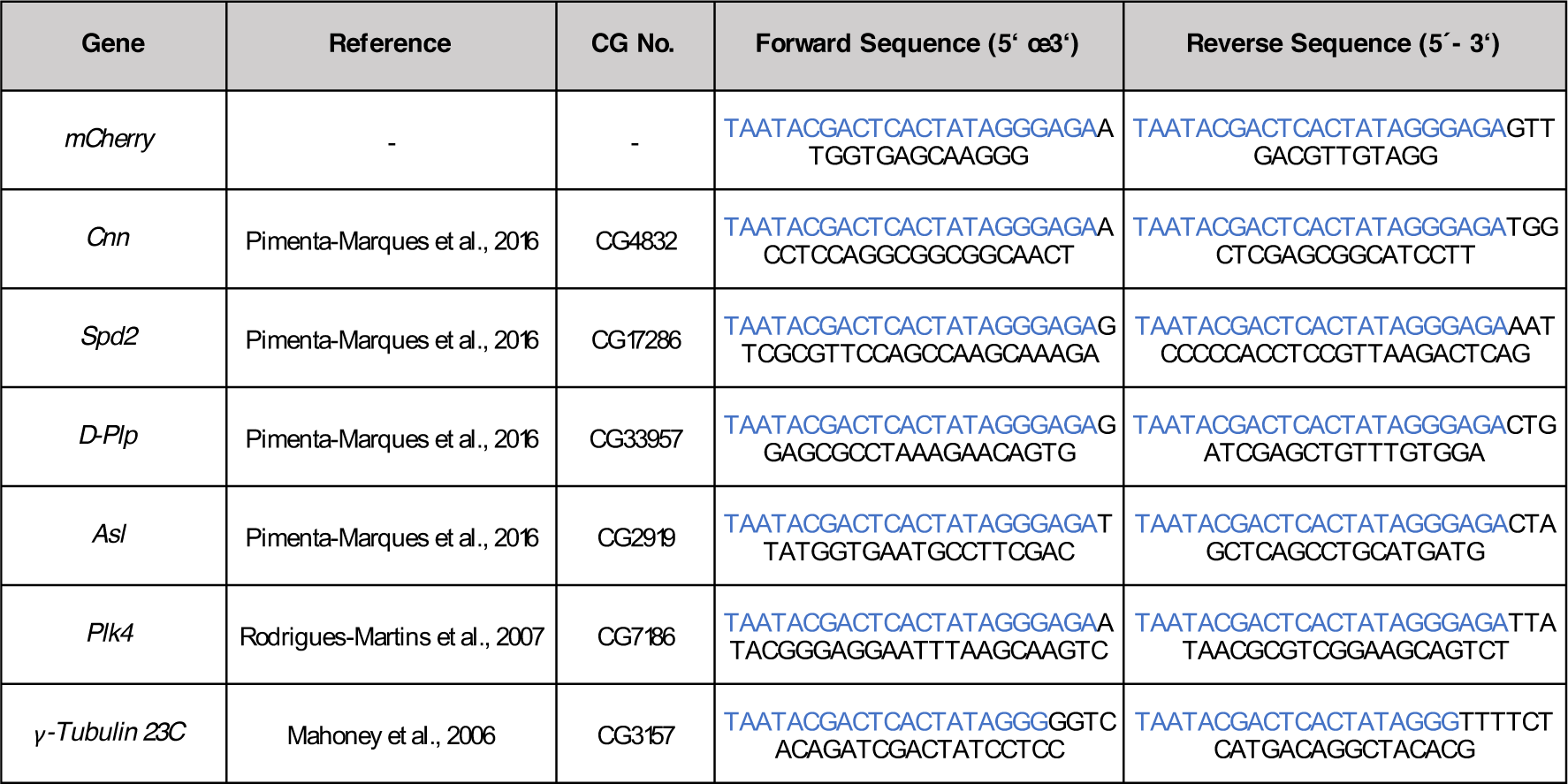
List of primers used for dsRNA synthesis. The overhangs for in vitro transcription with the T7 RNA polymerase are depicted in blue.

**Supplementary Table 6.**
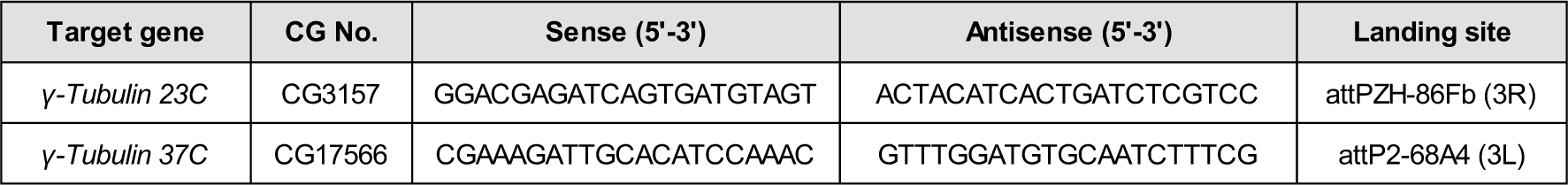
Sequences of the oligonucleotides used to generate short hairpin RNA (shRNA) targeting different *Drosophila melanogaster* gene products. Each combination of oligos was annealed and cloned into pWALIUM22, to drive knock-down of each target gene specifically in the female germline.

**Supplementary Table 7.**
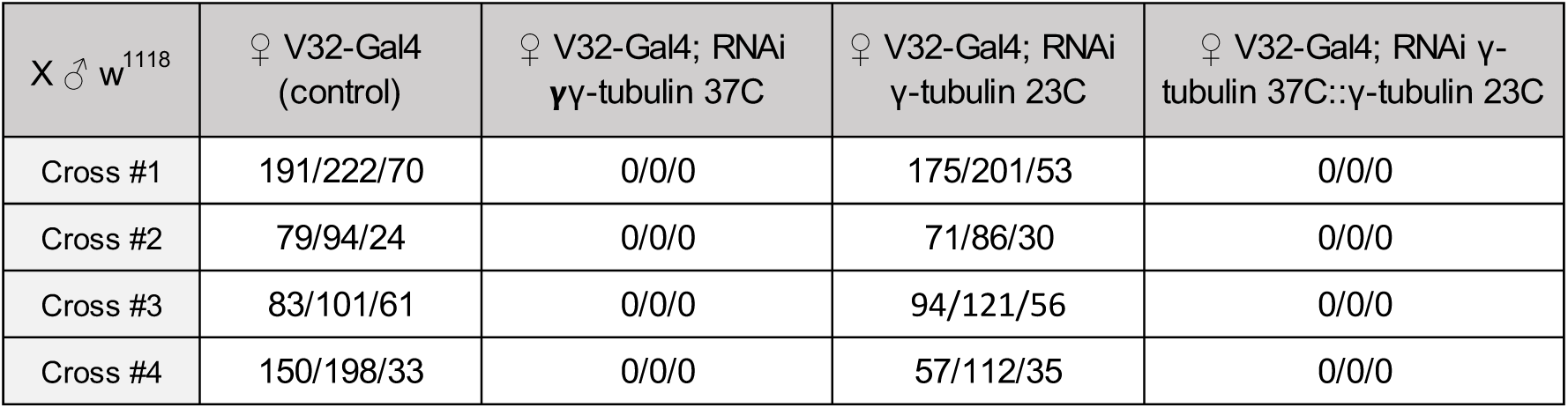
Lethality assay to determine viability of the shRNA fly lines. Number of pupae per vial in crosses between females carrying the V32-Gal4 and shRNA against γ-tubulin 37C and/or γ-tubulin 23C and w1118 males. V32-Gal4 females were crossed to w1118 males as control. For each genotype, four independent crosses were performed, with three technical repeats.

| X ♂ w<sup>1118</sup> | ♀ V32-Gal4 (control) | ♀ V32-Gal4; RNAi γγ-tubulin 37C | ♀ V32-Gal4; RNAi γ-tubulin 23C | ♀ V32-Gal4; RNAi γ-tubulin 37C::γ-tubulin 23C |
| --- | --- | --- | --- | --- |
| Cross #1 | 191/222/70 | 0/0/0 | 175/201/53 | 0/0/0 |
| Cross #2 | 79/94/24 | 0/0/0 | 71/86/30 | 0/0/0 |
| Cross #3 | 83/101/61 | 0/0/0 | 94/121/56 | 0/0/0 |
| Cross #4 | 150/198/33 | 0/0/0 | 57/112/35 | 0/0/0 |

## SUPPLEMENTARY TIME-LAPSE MOVIES

**Movie 1. (In support of Figure 1) - Centriole biogenesis in a *Drosophila melanogaster* egg explant.** Maximum intensity Z-projection from a time-lapse movie of a cytosolic explant isolated from an unfertilised Drosophila egg overexpressing Plk4, acquired on a spinning-disk confocal microscope. Centrioles are absent in the first time point and form de novo throughout the experiment detected as spots (Spd2, in green) associated with microtubule asters (magenta), reported by the microtubule associated protein Jupiter. Time (min:sec) is shown at the top left.

**Movies 2A–D (In support of Figure 2): Centrioles assemble de novo, recruit different centrosomal molecules and duplicate.** Maximum intensity Z-projection from time-lapse movies of droplets of cytosolic extract from non-cycling unfertilised Drosophila eggs overexpressing Plk4, acquired on a spinning-disk confocal microscope. Movies show centriole biogenesis reported by different centrosomal proteins in green – Plk4 (A), Ana1 (B), Asl (C) and Spd2 (D) – and the microtubule-associated protein Jupiter (magenta). The larger green blobs result from yolk autofluorescence, highly noticeable in the Plk4 movie. Time (min:sec) is shown at the top left of each four movies.

**Movie 3. (In support of Figure 5) mNeonGreen-Plk4 localisation in a syncytial *Drosophila* embryo.** Time-lapse movie of an embryo expressing homozygous mNeonGreen-Plk4 (endogenously labeled by CRISPR, in green) and RFP-Tubulin (magenta), acquired on a spinning-disk confocal microscope, through nuclear cycles 10-13. The movie is a bleach-corrected maximum intensity Z-projection. Time (min:sec) is shown at the top left.

## References

Aldrich, H.C. 1967. The Ultrastructure of Meiosis in Three Species of Physarum. Mycologia. 59:127– 148. doi:10.2307/3756947.

Aydogan, M.G., T.L. Steinacker, M. Mofatteh, L. Gartenmann, A. Wainman, S. Saurya, P.T. Conduit, F.Y. Zhou, M.A. Boemo, and J.W. Raff. 2019. A free-running oscillator times and executes centriole biogenesis. BioRxiv.

Banterle, N., and P. Gönczy. 2017. Centriole Biogenesis: From Identifying the Characters to Understanding the Plot. Annu. Rev. Cell Dev Biol. 33:23–49. doi: 10.1146/annurev-cellbio-100616-060454.

Bauer, M., F. Cubizolles, A. Schmidt, and E.A. Nigg. 2016. Quantitative analysis of human centrosome architecture by targeted proteomics and fluorescence imaging. EMBO J. 35:2152–2166. doi:10.15252/embj.

Bettencourt-Dias, M., A. Rodrigues-Martins, L. Carpenter, M. Riparbelli, L. Lehmann, M.K. Gatt, N. Carmo, F. Balloux, G. Callaini, and D.M. Glover. 2005. SAK/PLK4 is required for centriole duplication and flagella development. Curr. Biol. 15:2199–2207. doi:10.1016/j.cub.2005.11.042.

Bettencourt-Dias, M., F. Hildebrandt, D. Pellman, G. Woods, and S.A. Godinho. 2011. Centrosomes and cilia in human disease. Trends Genet. 27:307–15. doi:10.1016/j.tig.2011.05.004.

Bettencourt-Dias, M., R. Giet, R. Sinka, A. Mazumdar, W.G. Lock, F. Balloux, P.J. Zafiropoulos, S. Yamaguchi, S. Winter, R.W. Carthew, M. Cooper, D. Jones, L. Frenz, and D.M. Glover. 2004. Genome-wide survey of protein kinases required for cell cycle progression. Nature. 432:980–987. doi.org/10.1038/nature03160.

Blachon, S., J. Gopalakrishnan, Y. Omori, A. Polyanovsky, A. Church, D. Nicastro, J. Malicki, and T. Avidor-Reiss. 2008. Drosophila asterless and vertebrate Cep152 Are orthologs essential for centriole duplication. Genetics. 180:2081–94. doi:10.1534/genetics.108.095141.

Boese, C.J., J. Nye, D.W. Buster, T.A. Mclamarrah, A.E. Byrnes, K.C. Slep, N.M. Rusan, and G.C. Rogers. 2018. Asterless is a Polo-like kinase 4 substrate that both activates and inhibits kinase activity depending on its phosphorylation state. Mol. Biol. Cell. 29:2874–2886. doi: 10.1091/mbc.

Breslow, D.K., and A.J. Holland. 2019. Mechanism and Regulation of Centriole and Cilium Biogenesis. Annu. Rev. Biochem. 88:11.1-11.34. doi: 10.1146/annurev-biochem-013118-111153.

Chang, C., W. Hsu, J. Tsai, C.C. Tang, and T.K. Tang. 2016. CEP295 interacts with microtubules and is required for centriole elongation. J. Cell Sci. 129:2501–2513. doi: 10.1242/jcs.186338.

Charvin, G., C. Oikonomou, E.D. Siggia, and F.R. Cross. 2009. Origin of Irreversibility of Cell Cycle Start in Budding Yeast. PLoS Biol. 8. doi:10.1371/journal.pbio.1000284.

Citron, Y.R., C.J. Fagerstrom, B. Keszthelyi, B. Huang, N.M. Rusan, M.J.S. Kelly, and D.A. Agard. 2018. The centrosomin CM2 domain is a multifunctional binding domain with distinct cell cycle roles. PLoS One. 13:1–15. doi:10.1371/journal.pone.0190530.

Cizmecioglu, O., M. Arnold, R. Bahtz, F. Settele, L. Ehret, U. Haselmann-Weiss, C. Antony, and I. Hoffmann. 2010. Cep152 acts as a scaffold for recruitment of Plk4 and CPAP to the centrosome. J. Cell Biol. 191:731–9. doi:10.1083/jcb.201007107.

Conduit, P. T., Richens, J. H., Wainman, A., Holder, J., Vicente, C. C., Pratt, M. B., Dix, C. I., Novak, Z. a, Dobbie, I. M., Schermelleh, L., and J.W. Raff (2014). A molecular mechanism of mitotic centrosome assembly in Drosophila. Elife 3:e03399. doi: 10.7554/eLife.03399.

Conduit, P.T., A. Wainman, Z.A. Novak, T.T. Weil, and J.W. Raff. 2015. Re-examining the role of Drosophila Sas-4 in centrosome assembly using two-colour-3D-SIM FRAP. Elife. 4. doi:10.7554/eLife.08483.

Courtois, A., M. Schuh, J. Ellenberg, and T. Hiiragi. 2012. The transition from meiotic to mitotic spindle assembly is gradual during early mammalian development. J. Cell Biol. 198:357–370. doi:10.1083/jcb.201202135.

Cunha-Ferreira, I., A. Rodrigues-Martins, I. Bento, M. Riparbelli, W. Zhang, E. Laue, G. Callaini, D.M. Glover, and M. Bettencourt-Dias. 2009. The SCF/Slimb ubiquitin ligase limits centrosome amplification through degradation of SAK/PLK4. Curr. Biol. 19:43–49. doi:10.1016/j.cub.2008.11.037.

Cunha-Ferreira, I., I. Bento, A. Pimenta-Marques, S.C. Jana, M. Lince-Faria, P. Duarte, J. Borrego-Pinto, S. Gilberto, T. Amado, D. Brito, A. Rodrigues-Martins, J. Debski, N. Dzhindzhev, and M. Bettencourt-Dias. 2013. Regulation of Autophosphorylation Controls PLK4 Self-Destruction and Centriole Number. Curr. Biol. 23:2245–54. doi:10.1016/j.cub.2013.09.037.

Dammermann, A., T. Müller-Reichert, L. Pelletier, B. Habermann, A. Desai, and K. Oegema. 2004. Centriole assembly requires both centriolar and pericentriolar material proteins. Dev. Cell. 7:815–829. doi:10.1016/j.devcel.2004.10.015.

de-Carvalho, J., O. Deshpande, C. Nabais, and I.A. Telley. 2018. A cell-free system of Drosophila egg explants supporting native mitotic cycles. 144. 1st ed. Elsevier Inc. 233–257 pp. doi: 10.1016/bs.mcb.2018.03.011.

De-Carvalho, J., S. Tlili, L. Hufnagel, T. Saunders, and I. Telley. 2020. Aster repulsion drives local ordering in an active system. bioRxiv. doi:10.1101/2020.06.04.133579.

Deneke, V.E., A. Puliafito, D. Krueger, A. V Narla, A. De Simone, L. Primo, M. Vergassola, S. De Renzis, and S. Di Talia. 2019. Self-Organized Nuclear Positioning Synchronizes the Cell Cycle in Drosophila Embryos. Cell. 177:925–941. doi:10.1016/j.cell.2019.03.007.

Dirksen, E.R. 1961. The presence of centrioles in artificially activated sea urchin eggs. J. Cell Biol. 11:244–247. doi: 10.1083/jcb.11.1.244.

Dzhindzhev, N.S., Q.D. Yu, K. Weiskopf, G. Tzolovsky, I. Cunha-Ferreira, M. Riparbelli, A. Rodrigues-Martins, M. Bettencourt-Dias, G. Callaini, and D.M. Glover. 2010. Asterless is a scaffold for the onset of centriole assembly. Nature. 467:714–8. doi:10.1038/nature09445.

Feng, Z., A. Caballe, A. Wainman, S. Johnson, A.F.M. Haensele, M.A. Cottee, P.T. Conduit, L.M. Susan, and J.W. Raff. 2017. Structural Basis for Mitotic Centrosome Assembly in Flies. Cell. 169:1078–1089. doi:10.1016/j.cell.2017.05.030.

Ferree, P.M., K. McDonald, B. Fasulo, and W. Sullivan. 2006. The origin of centrosomes in parthenogenetic hymenopteran insects. Curr. Biol. 16:801–7. doi:10.1016/j.cub.2006.03.066.

Fu, J., Z. Lipinszki, H. Rangone, M. Min, C. Mykura, J. Chao-chu, S. Schneider, N.S. Dzhindzhev, M. Gottardo, G. Riparbelli, G. Callaini, and D.M. Glover. 2016. Conserved Molecular Interactions in Centriole-to-Centrosome Conversion. Nat. Cell Biol. 18:87–99. doi:10.1038/ncb3274.

Godinho, S.A., and D. Pellman. 2014. Causes and consequences of centrosome abnormalities in cancer. Philos. Trans. R. Soc. B Biol. Sci. 369. doi:10.1098/rstb.2013.0467.

Godinho, S.A., R. Picone, M. Burute, R. Dagher, Y. Su, C.T. Leung, K. Polyak, J.S. Brugge, M. Thery, and D. Pellman. 2014. Oncogene-like induction of cellular invasion from centrosome amplification. Nature. 510:167–171. doi:10.1038/nature13277.

Gomez-Ferreria, M.A., U. Rath, D.W. Buster, S.K. Chanda, J.S. Caldwell, D.R. Rines, and D.J. Sharp. 2007. Human Cep192 Is Required for Mitotic Centrosome and Spindle Assembly. Curr. Biol. 17:1960– 1966. doi:10.1016/j.cub.2007.10.019.

Grimes, G.W. 1973a. Origin and development of kinetosomes in Oxytricha fallax. J. Cell Sci. 13:43–53.

Grimes, G.W. 1973b. Morphological discontinuity of kinetosomes during the life cycle of Oxytricha fallax. J. Cell Biol. 57:229–232. doi: 10.1083/jcb.57.1.229.

Guderian, G., J. Westendorf, A. Uldschmid, and E. A. Nigg. 2010. Plk4 trans-autophosphorylation regulates centriole number by controlling betaTrCP-mediated degradation. J. Cell Sci. 123:2163–2169. doi:10.1242/jcs.068502.

Gueth-Hallonet, C., C. Antony, J. Aghion, A. Santa-Maria, I. Lajoie-Mazenc, M. Wright, and B. Maro. 1993. γ-Tubulin is present in acentriolar MTOCs during early mouse development. J. Cell Sci. 105:157– 166.

Habedanck, R., Y.D. Stierhof, C.J. Wilkinson, and E.A. Nigg. 2005. The Polo kinase PLK4 functions in centriole duplication. Nat. Cell Biol. 7:1140–1146. doi: 10.1038/ncb1320.

Harvey, E.B. 1936. Parthenogenetic Merogony or Cleavage without Nuclei in Arbacia punctulata. Mar. Biol. Lab. 71:101–121. doi: 10.2307/1537411.

Holland, A.J., D. Fachinetti, Q. Zhu, M. Bauer, I.M. Verma, E. A. Nigg, and D.W. Cleveland. 2012. The autoregulated instability of Polo-like kinase 4 limits centrosome duplication to once per cell cycle. Genes Dev. 26:2684–2689. doi:10.1101/gad.207027.112.

Horner, V.L., A. Czank, J.K. Jang, N. Singh, B.C. Williams, J. Puro, E. Kubli, S.D. Hanes, K.S. McKim, M.F. Wolfner, and M.L. Goldberg. 2006. The Drosophila Calcipressin Sarah Is Required for Several Aspects of Egg Activation. Curr. Biol. 16:1441–1446. doi:10.1016/j.cub.2006.06.024.

Idei, M., K. Osada, S. Sato, T. Nakayama, T. Nagumo, and D.G. Mann. 2013. Sperm ultrastructure in the diatoms Melosira and Thalassiosira and the significance of the 9 + 0 configuration. Protoplasma. 250:833–850. doi:10.1007/s00709-012-0465-8.

Ito, D., S. Zitouni, S.C. Jana, P. Duarte, J. Surkont, Z. Carvalho-Santos, J.B. Pereira-Leal, M.G. Ferreira, and M. Bettencourt-Dias. 2019. Pericentrin-mediated SAS-6 recruitment promotes centriole assembly. Elife. 8. doi: 10.7554/eLife.41418.

Izquierdo, D., W.J. Wang, K. Uryu, and M.F.B. Tsou. 2014. Stabilization of cartwheel-less centrioles for duplication requires CEP295-mediated centriole to centrosome conversion. Cell. Rep. 8:957–965. doi:10.1016/j.celrep.2014.07.022.

Jaqaman, K., D. Loerke, M. Mettlen, H. Kuwata, S. Grinstein, S.L. Schmid, and G. Danuser. 2008. Robust single-particle tracking in live-cell time-lapse sequences. Nat. Methods. 5:695–702. doi:10.1038/nmeth.1237.

Joukov, V., and A. De Nicolo. 2019. The Centrosome and the Primary Cilium: The Yin and Yang of a Hybrid Organelle. Cells. 8. doi: 10.3390/cells8070701.

Keller, D., M. Orpinell, N. Olivier, M. Wachsmuth, R. Mahen, R. Wyss, V. Hachet, J. Ellenberg, S. Manley, and P. Gönczy. 2014. Mechanisms of HsSAS-6 assembly promoting centriole formation in human cells. J. Cell Biol. 204:697–712. doi:10.1083/jcb.201307049.

Khodjakov, A., C.L. Rieder, G. Sluder, G. Cassels, O. Sibon, and C. Wang. 2002. De novo formation of centrosomes in vertebrate cells arrested during S phase. J. Cell Biol. 158:1171–1181. doi:10.1083/jcb.200205102.

Kitazawa, D., T. Matsuo, K. Kaizuka, C. Miyauchi, D. Hayashi, and Y.H. Inoue. 2014. Orbit/CLASP is required for myosin accumulation at the cleavage furrow in Drosophila male meiosis. PLoS One. 9. doi:10.1371/journal.pone.0093669.

Klebba, J.E., B.J. Galletta, J. Nye, K.M. Plevock, D.W. Buster, N. A Hollingsworth, K.C. Slep, N.M. Rusan, and G.C. Rogers. 2015b. Two Polo-like kinase 4 binding domains in Asterless perform distinct roles in regulating kinase stability. J. Cell Biol. 208:401–414. doi:10.1083/jcb.201410105.

Klebba, J.E., D.W. Buster, A.L. Nguyen, S. Swatkoski, M. Gucek, N.M. Rusan, and G.C. Rogers. 2013. Polo-like Kinase 4 Autodestructs by Generating Its Slimb-Binding Phosphodegron. Curr. Biol. 23:2255– 61. doi:10.1016/j.cub.2013.09.019.

Klebba, J.E., D.W. Buster, T.A. McLamarrah, N.M. Rusan, and G.C. Rogers. 2015a. Autoinhibition and relief mechanism for Polo-like kinase 4. Proc. Natl. Acad. Sci. 112:E657–E666. doi:10.1073/pnas.1417967112.

Kleylein-Sohn, J., J. Westendorf, M. Le Clech, R. Habedanck, Y.D. Stierhof, and E. A Nigg. 2007. Plk4-induced centriole biogenesis in human cells. Dev. Cell. 13:190–202. doi:10.1016/j.devcel.2007.07.002.

La Terra, S., C.N. English, P. Hergert, B.F. McEwen, G. Sluder, and A. Khodjakov. 2005. The de novo centriole assembly pathway in HeLa cells: cell cycle progression and centriole assembly/maturation. J. Cell Biol. 168:713–722. doi:10.1083/jcb.200411126.

Lambrus, B., K.M. Clutario, V. Daggubati, M. Snyder, G. Sluder, and A. Holland. 2015. p53 protects against genome instability following centriole duplication failure. J. Cell Biol. 210:63–77. doi: 10.1083/jcb.201502089.

Leda, M., A.J. Holland, and A.B. Goryachev. 2018. Autoamplification and Competition Drive Symmetry Breaking: Initiation of Centriole Duplication by the PLK4-STIL Network. iScience. 8:222–235. doi:10.1016/j.isci.2018.10.003.

Lerit, D.A., H.A. Jordan, J.S. Poulton, C.J. Fagerstrom, B.J. Galletta, M. Peifer, and N.M. Rusan. 2015. Interphase centrosome organization by the PLP-Cnn scaffold is required for centrosome function. J. Cell Biol. 210:79–97. doi:10.1083/jcb.201503117.

Levine, M.S., B. Bakker, B. Boeckx, J. Moyett, J. Lu, D.C. Spierings, P.M. Lansdorp, D.W. Cleveland, F. Foijer, and A.J. Holland. 2018. Centrosome amplification is sufficient to promote spontaneous tumorigenesis in mammals. Dev. Cell. 40:313–322. doi:10.1016/j.devcel.2016.12.022.

Loncarek, J., P. Hergert, V. Magidson, and A. Khodjakov. 2008. Control of daughter centriole formation by the pericentriolar material. Nat. Cell Biol. 10:322–328.

Loncarek, J., and A. Khodjakov. 2009. Ab ovo or de novo? Mechanisms of centriole duplication. Mol. Cells. 27:135–142. doi:10.1007/s10059-009-0017-z.

Lopes, C.A.M., M. Mesquita, A.I. Cunha, J. Cardoso, S. Carapeta, C. Laranjeira, A.E. Pinto, J.B. Pereira-Leal, A. Dias-Pereira, M. Bettencourt-Dias, and P. Chaves. 2018. Centrosome amplification arises before neoplasia and increases upon p53 loss in tumorigenesis. J. Cell Biol. 217:2353–2363. doi:10.1083/jcb.201711191.

Lopes, C.A.M., S.C. Jana, I. Cunha-Ferreira, S. Zitouni, I. Bento, P. Duarte, S. Gilberto, F. Freixo, A. Guerrero, M. Francia, M. Lince-Faria, J. Carneiro, and M. Bettencourt-Dias. 2015. PLK4 trans-Autoactivation Controls Centriole Biogenesis in Space. Dev. Cell. 35:222–235. doi:10.1016/j.devcel.2015.09.020.

Lowe, N., J.S. Rees, J. Roote, E. Ryder, I.M. Armean, G. Johnson, E. Drummond, H. Spriggs, J. Drummond, J.P. Magbanua, H. Naylor, R. Bastock, S. Huelsmann, V. Trovisco, M. Landgraf, S. Knowles-barley, J.D. Armstrong, H. White-cooper, C. Hansen, R.G. Phillips, T. Uk, D. Protein, S. Consortium, K.S. Lilley, S. Russell, and D.S. Johnston. 2014. Analysis of the expression patterns, subcellular localisations and interaction partners of Drosophila proteins using a pigP protein trap library. Development. 141:3994–4005. doi:10.1242/dev.111054.

Mahen, R., A.D. Jeyasekharan, N.P. Barry, and A.R. Venkitaraman. 2011. Continuous polo-like kinase 1 activity regulates diffusion to maintain centrosome self-organization during mitosis. Proc. Natl. Acad. Sci. 108:9310–9315. doi: 10.1073/pnas.1101112108.

Mahoney, N.M., G. Goshima, A.D. Douglass, and R.D. Vale. 2006. Making Microtubules and Mitotic Spindles in Cells without Functional Centrosomes. Curr. Biol. 564–569. doi:10.1016/j.cub.2006.01.053.

Majer, G., and K. Zick. 2015. Accurate and absolute diffusion measurements of Rhodamine 6G in low-concentration aqueous solutions by the PGSE-WATERGATE sequence. J. Chem. Phys. 142. doi: 10.1063/1.4919054.

Marshall, W.F., Y. Vucica, and J.L. Rosenbaum. 2001. Kinetics and regulation of de novo centriole assembly: Implications for the mechanism of centriole duplication. Curr. Biol. 11:308–17. doi: 10.1016/s0960-9822(01)00094-x.

Marteil, G., A. Guerrero, A.F. Vieira, B.P. De Almeida, P. Machado, S. Mendonça, M. Mesquita, B. Villarreal, I. Fonseca, M.E. Francia, K. Dores, N.P. Martins, S.C. Jana, E.M. Tranfield, N.L. Barbosa-Morais, J. Paredes, D. Pellman, S.A. Godinho, and M. Bettencourt-Dias. 2018. Over-elongation of centrioles in cancer promotes centriole amplification and chromosome missegregation. Nat. Commun. 9. doi:10.1038/s41467-018-03641-x.

Mclamarrah, T.A., D.W. Buster, B.J. Galletta, C.J. Boese, J.M. Ryniawec, N.A. Hollingsworth, A.E. Byrnes, C.W. Brownlee, K.C. Slep, N.M. Rusan, and G.C. Rogers. 2018. An ordered pattern of Ana2 phosphorylation by Plk4 is required for centriole assembly. J. Cell Biol. 217:1217–1231. doi: 10.1083/jcb.201605106.

Mercey, O., A.A. Jord, P. Rostaing, A. Mahuzier, A. Fortoul, A. Boudjema, M. Faucourt, N. Spassky, and A. Meunier. 2019a. Dynamics of centriole amplification in centrosome-depleted brain multiciliated progenitors. Sci. Rep. 9. doi:10.1038/s41598-019-49416-2.

Mercey, O., M.S. Levine, G.M. Lomastro, P. Rostaing, E. Brotslaw, V. Gomez, A. Kumar, N. Spassky, B.J. Mitchell, A. Meunier, and A.J. Holland. 2019b. Massive centriole production can occur in the absence of deuterosomes in multiciliated cells. Nat. Cell Biol. 12:1544–1552. doi:10.1038/s41556-019-0427-x.

Mir, L., M. Wright, and A. Moisand. 1984. Variations in the number of centrioles, the number of microtubule organizing centers 1 and the percentage of mitotic abnormalities in Physarum polycephalum amoebae. Protoplasma. 120:20–35. doi: 10.1007/BF01287614.

Mizukami, I., and J. Gall. 1966. Centriole replication. II. Sperm formation in the fern, Marsilea, and the cycad, Zamia. J. Cell Biol. 29:97–111. doi:10.1083/jcb.29.1.97.

Montenegro Gouveia, S., S. Zitouni, D. Kong, P. Duarte, B. Ferreira Gomes, A.L. Sousa, E.M. Tranfield, A. Hyman, J. Loncarek, and M. Bettencourt-Dias. 2018. PLK4 is a microtubule-associated protein that self-assembles promoting de novo MTOC formation. J. Cell Sci. 132. doi:10.1242/jcs.219501.

Moyer, T.C., K.M. Clutario, B.G. Lambrus, V. Daggubati, and A.J. Holland. 2015. Binding of STIL to Plk4 activates kinase activity to promote centriole assembly. J. Cell Biol. 209:863–878. doi:10.1083/jcb.201502088.

Nabais, C., S.G. Pereira, and M. Bettencourt-Dias. 2018. Noncanonical Biogenesis of Centrioles and Basal Bodies. Cold Spring Harb. Symp. Quant. Biol. 82:123–135. doi:10.1101/sqb.2017.82.034694.

Park, J.E., L. Zhang, J.K. Bang, T. Andresson, F. Di Maio, and K.S. Lee. 2019. Phase separation of Polo-like kinase 4 by autoactivation and clustering drives centriole biogenesis. Nat. Commun. 10. doi:10.1038/s41467-019-12619-2.

Peel, N., N.R. Stevens, R. Basto, and J.W. Raff. 2007. Overexpressing centriole-replication proteins in vivo induces centriole overduplication and de novo formation. Curr. Biol. 17:834–843. doi:10.1016/j.cub.2007.04.036.

Pimenta-Marques, A., I. Bento, C.A.M. Lopes, P. Duarte, S.C. Jana, and M. Bettencourt-Dias. 2016. A mechanism for the elimination of the female gamete centrosome in Drosophila melanogaster. Science. 353:aaf4866. doi:10.1126/science.aaf4866.

Port, F., H.M. Chen, T. Lee, and S.L. Bullock. 2014. Optimized CRISPR/Cas tools for efficient germline and somatic genome engineering in Drosophila. Proc. Natl. Acad. Sci. U. S. A. doi:10.1073/pnas.1405500111.

Prudêncio, P., and L.G. Guilgur. 2015. Protein Extraction from Drosophila Embryos and Ovaries. Bio-protocol. 5:e1459. doi:10.21769/BioProtoc.1459.

Rale, M.J., R.S. Kadzik, and S. Petry. 2018. Phase Transitioning the Centrosome into a Microtubule Nucleator. Biochemistry. 57:30–37. doi:10.1021/acs.biochem.7b01064.

Renzaglia, K.S., and D.J. Garbary. 2001. Motile Gametes of Land Plants: Diversity, Development, and Evolution. CRC. Crit. Rev. Plant Sci. 20:107–213. doi:10.1080/20013591099209.

Riparbelli, M.G., and G. Callaini. 2003. Drosophila parthenogenesis: A model for de novo centrosome assembly. Dev. Biol. 260:298–313. doi:10.1016/S0012-1606(03)00243-4.

Riparbelli, M.G., and G. Callaini. 2005. The meiotic spindle of the Drosophila oocyte: the role of Centrosomin and the central aster. J. Cell Sci. 118:2827–2836. doi:10.1242/jcs.02413.

Riparbelli, M.G., D. Tagu, J. Bonhomme, and G. Callaini. 2005. Aster self-organization at meiosis: a conserved mechanism in insect parthenogenesis ? Dev. Biol. 278:220–230. doi:10.1016/j.ydbio.2004.11.009.

Riparbelli, M.G., R. Stouthamer, R. Dallai, and G. Callaini. 1998. Microtubule Organization during the Early Development of the Parthenogenetic Egg of the Hymenopteran Muscidifurax uniraptor. Dev. Biol. 195:89–99. doi:10.1006/dbio.1997.8841.

Rodrigues-martins, A., M. Riparbelli, G. Callaini, D.M. Glover, and M. Bettencourt-Dias. 2008. From centriole biogenesis to cellular function: Centrioles are essential for cell division at critical developmental stages. Cell Cycle. 7:11–16. doi:10.4161/cc.7.1.5226.

Rodrigues-Martins, A., M. Riparbelli, G. Callaini, D.M. Glover, and M. Bettencourt-Dias. 2007. Revisiting the role of the mother centriole in centriole biogenesis. Science. 316:1046–50. doi:10.1126/science.1142950.

Schindelin, J., I. Arganda-Carreras, E. Frise, V. Kaynig, M. Longair, T. Pietzsch, S. Preibisch, C. Rueden, S. Saalfeld, B. Schmid, J.Y. Tinevez, D.J. White, V. Hartenstein, K. Eliceiri, P. Tomancak1, and A. Cardona. 2012. Fiji - an Open Source platform for biological image analysis Johannes. Nat. Methods. 9. doi:10.1038/nmeth.2019.Fiji.

Shohei, Y., and D. Kitagawa. 2018. Self-organization of Plk4 regulates symmetry breaking in centriole duplication. BioRxiv.

Takao, D., S. Yamamoto, and D. Kitagawa. 2019. A theory of centriole duplication based on self-organized spatial pattern formation. J. Cell Biol. 218:3537–3547.doi: 10.1083/jcb.201904156.

Tavosanis, G., S. Llamazares, G. Goulielmos, and C. Gonzalez. 1997. Essential role for γ -tubulin in the acentriolar female meiotic spindle of Drosophila. EMBO J. 16:1809–1819. doi: 10.1093/emboj/16.8.1809.

Telley, I. a, I. Gáspár, A. Ephrussi, and T. Surrey. 2013. A single Drosophila embryo extract for the study of mitosis ex vivo. Nat. Protoc. 8:310–24. doi:10.1038/nprot.2013.003.

Telley, I.A., I. Gáspár, A. Ephrussi, and T. Surrey. 2012. Aster migration determines the length scale of nuclear separation in the Drosophila syncytial embryo. J. Cell Biol. 197:887–895. doi:10.1083/jcb.201204019.

Tram, U., and W. Sullivan. 2000. Reciprocal inheritance of centrosomes in the parthenogenetic Hymenopteran Nasonia vitripennis. Curr. Biol. 10:1413–1419. doi:10.1016/S0960-9822(00)00795-8.

Tsuchiya, Y., S. Yoshiba, A. Gupta, K. Watanabe, and D. Kitagawa. 2016. Cep295 is a conserved scaffold protein required for generation of a bona fide mother centriole. Nat. Commun. 7:1–13. doi:10.1038/ncomms12567.

Tyson, J.J., and B. Novak. 2001. Regulation of the Eukaryotic Cell Cycle: Molecular Antagonism, Hysteresis, and Irreversible Transitions. J. Theor. Biol. 210:249–263. doi:10.1006/jtbi.2001.2293.

Uetake, Y., J. Loncarek, J.J. Nordberg, C.N. English, S. La Terra, A. Khodjakov, and G. Sluder. 2007. Cell cycle progression and de novo centriole assembly after centrosomal removal in untransformed human cells. J. Cell Biol. 176:173–182. doi:10.1083/jcb.200607073.

Vardy, L., and T.L. Orr-Weaver. 2007. The Drosophila PNG Kinase Complex Regulates the Translation of Cyclin B. Dev. Cell. 12:157–166. doi:10.1016/j.devcel.2006.10.017.

Varmark, H., S. Llamazares, E. Rebollo, B. Lange, J. Reina, H. Schwarz, and C. Gonzalez. 2007. Asterless Is a Centriolar Protein Required for Centrosome Function and Embryo Development in Drosophila. Curr. Biol. 17:1735–1745. doi:10.1016/j.cub.2007.09.031.

Wang, W.J., R.K. Soni, K. Uryu, and M.F.B. Tsou. 2011. The conversion of centrioles to centrosomes: essential coupling of duplication with segregation. J. Cell Biol. 193:727–739. doi:10.1083/jcb.201101109.

Wong, Y.L., J. V Anzola, R.L. Davis, M. Yoon, A. Motamedi, A. Kroll, C.P. Seo, J.E. Hsia, S.K. Kim, J.W. Mitchell, J. Brian, A. Desai, T.C. Gahman, A.K. Shiau, and K. Oegema. 2015. Reversible centriole depletion with an inhibitor of Polo-like kinase 4. Science. 348:1155–1160. doi:10.1126/science.aaa5111.

Yamamoto, S., and D. Kitagawa. 2019. Self-organization of Plk4 regulates symmetry breaking in centriole duplication. Nat. Commun. 10. doi:10.1038/s41467-019-09847-x.

Yatsu, N. 1905. The formation of centrosomes in enucleated egg-fragments. J. Exp. Zool. 2. doi:10.1002/jez.1400020206.

Zitouni, S., M.E. Francia, F. Leal, S.M. Gouveia, C. Nabais, P. Duarte, S. Gilberto, D. Brito, T. Moyer, M. Ohta, D. Kitagawa, A.J. Holland, E. Karsenti, T. Lorca, M. Lince-Faria, and M. Bettencourt-Dias. 2016. CDK1 prevents unscheduled PLK4-STIL complex assembly in centriole biogenesis. Curr. Biol. 26:1127–1137. doi:10.1016/j.cub.2016.03.055.

